# Programmable patterns in a DNA-based reaction-diffusion system

**DOI:** 10.1101/556381

**Authors:** Sifang Chen, Georg Seelig

## Abstract

Biology offers compelling proof that macroscopic “living materials” can emerge from reactions between diffusing biomolecules. Here, we show that molecular self-organization could be a similarly powerful approach for engineering functional synthetic materials. We introduce a programmable DNA-hydrogel that produces tunable patterns at the centimeter length scale. We generate these patterns by implementing chemical reaction networks through synthetic DNA complexes, embedding the complexes in hydrogel, and triggering with locally applied input DNA strands. We first demonstrate ring pattern formation around a circular input cavity and show that the ring width and intensity can be predictably tuned. Then, we create patterns of increasing complexity, including concentric rings and non-isotropic patterns. Finally, we show “destructive” and “constructive” interference patterns, by combining several ring-forming modules in the gel and triggering them from multiple sources. We further show that computer simulations based on the reaction-diffusion model can predict and inform the programming of target patterns.

## Introduction

Programmable matter research aims to engineer functional materials that can autonomously transform their appearances or physical properties in response to environmental stimuli and user-defined inputs. Top down methods like 3D-printing have enabled the development of shapeshifting biomimetic constructs that are sensitive to heat, light, or water^1, 2^. Advances in micro-robotics have led to modular robotic swarms that can self-organize into two- and three-dimensional structures^3, 4^. But we are still far from creating true programmable matter. Currently, synthetic materials and systems either rely on components too large to be integrated into material fabrics, as with modular robot systems, or have limited functions, as with 3D printed materials.

Chemical computing offers a tantalizing alternative. Biological patterning processes like camouflaging and morphogenesis suggest that complex and environmentally responsive systems could arise from the self-organization of information-bearing agents like molecules or cells^5, 6^. Engineering molecular systems to predictably form complex patterns like those seen in biology would clearly have significant implications for programmable materials research.

The mathematical model of reaction-diffusion provides a framework for designing and engineering programmable structures through chemical computing^6–8^. In this model, spatial patterns can emerge from local interactions between diffusing agents^9^. Simulations developed within this framework have successfully replicated complex biological patterns^10, 11^, suggesting a path toward model-guided engineering of autonomous self-organizing systems. However, experimental *de novo* realizations of pattern formation have been sparse.

Early examples of synthetic pattern formation include the Belousov-Zhabotinsky (BZ) chemical oscillator^12, 13^, which generates macroscopic spatiotemporal patterns via a series of redox reactions. While the mechanics of BZ reactions are well understood, we cannot control reaction kinetics or program the resulting patterns to display target behaviors. Synthetic biologists have genetically engineered quorum-sensing bacteria to create stripes and traveling waves^14–16^. Such results showcase the potential of a biochemical approach to programming self-organized pattern formation, but the precision of patterning is still limited because the engineered reaction networks operate in a background of evolved and not fully understood cellular machinery. Cell-free biochemical reaction networks are another promising alternative but systems engineered to date, although capable of generating a wide range of patterns in aqueous reactors^17–20^, still largely rely on catalysis of evolved enzymes and have limited programmability.

DNA is unique, even among biopolymers, in that interactions are quantitatively predictable and follow the rules of Watson-Crick base pairing^21, 22^. DNA origami and related self-assembly technologies take advantage of this predictability for the construction of 2- and 3-dimensional objects of varying sizes and complexity^23, 24^. This work has culminated in macroscopic materials with nanometer-scale addressability. But these periodic crystals^25^ or random gels^26, 27^ lack non-trivial long-range order and are expensive because DNA acts as the primary structural component. Thus, to recapitulate the diversity and scale of biological patterns and materials with DNA alone we still need to develop approaches that extend to the centimeter scale and beyond.

To address this need, recent work has begun to explore the feasibility of DNA-only reaction-diffusion patterns^28^. Toehold-mediated DNA strand displacement has proved to be a convenient framework for implementing complex reaction sequences using synthetic DNA in well-mixed test tubes^29, 30^. Using the principles of strand displacement, researchers have created sophisticated reaction networks that perform computation like neural networks^31, 32^, diagnostic classifiers^33^, dynamic 3D nanostructures^34, 35^ and even approximate the dynamics of formal, mathematically specified chemical reaction networks (CRNs)^36–39^. Building on these results, theoretical work has argued that a wide range of patterns is achievable if DNA-based CRNs are embedded in a spatial reactor^40, 41^. Chirieleison *et al.* took an important step toward experimentally demonstrating pattern formation with DNA strand displacement-based CRNs and engineered an edge detection system^42^. However, despite the advances made in these projects, the state of art for programming macroscopic features still lags that of their microscopic counterparts.

Here, we report the design and synthesis of a novel DNA-hydrogel hybrid material for programmable spatial patterning at the centimeter length scale. Patterns are generated via the reaction-diffusion of DNA complexes separately embedded in porous hydrogel and predefined cavities in the gel. Using this system, patterns of varying geometries can be generated and quantitatively tuned by controlling the reaction rates of species. To further demonstrate programmability, we show that the dynamic behavior of these spatial patterns can be predicted by computer simulations.

## Constructing a pulse-generator

Figure 1 shows the workflow of our DNA-based programmable patterning system. A simple ring-forming module forms the basic building block for all other patterns realized in this work (**Figure 1A**). Rings are an archetype for studying synthetic pattern formation^14, 16^ and, as we will show, form an ideal starting point for generating more complex patterns. To implement this module, we begin by formulating a pulse-generating CRN (**Fig. 1B**). We then realize this CRN using DNA strand displacement-based complexes (**Fig. 1C**). Next, we synthesize the DNA-hydrogel by suspending these DNA complexes in an agarose solution and molding the mixture into thin sheets. Finally, we load initiator strands into cavities in the DNA-hydrogel to trigger programmed pattern formation (**Fig. 1D**). A predictive spatial model built in Visual DSD informs the concentrations of initiator strands required for generating target patterns. Input parameters for the model include reaction constants and diffusion coefficients inferred from spectrometry and gel experiments (**Fig. 1D**). Depending on the initial conditions (concentrations of initiating strands) and boundary conditions (shape and placement of cavities), gels embedded with identical DNA gates can be programmed to display a variety of spatial dynamics (**Fig. 1E**).

**Figure 1.**
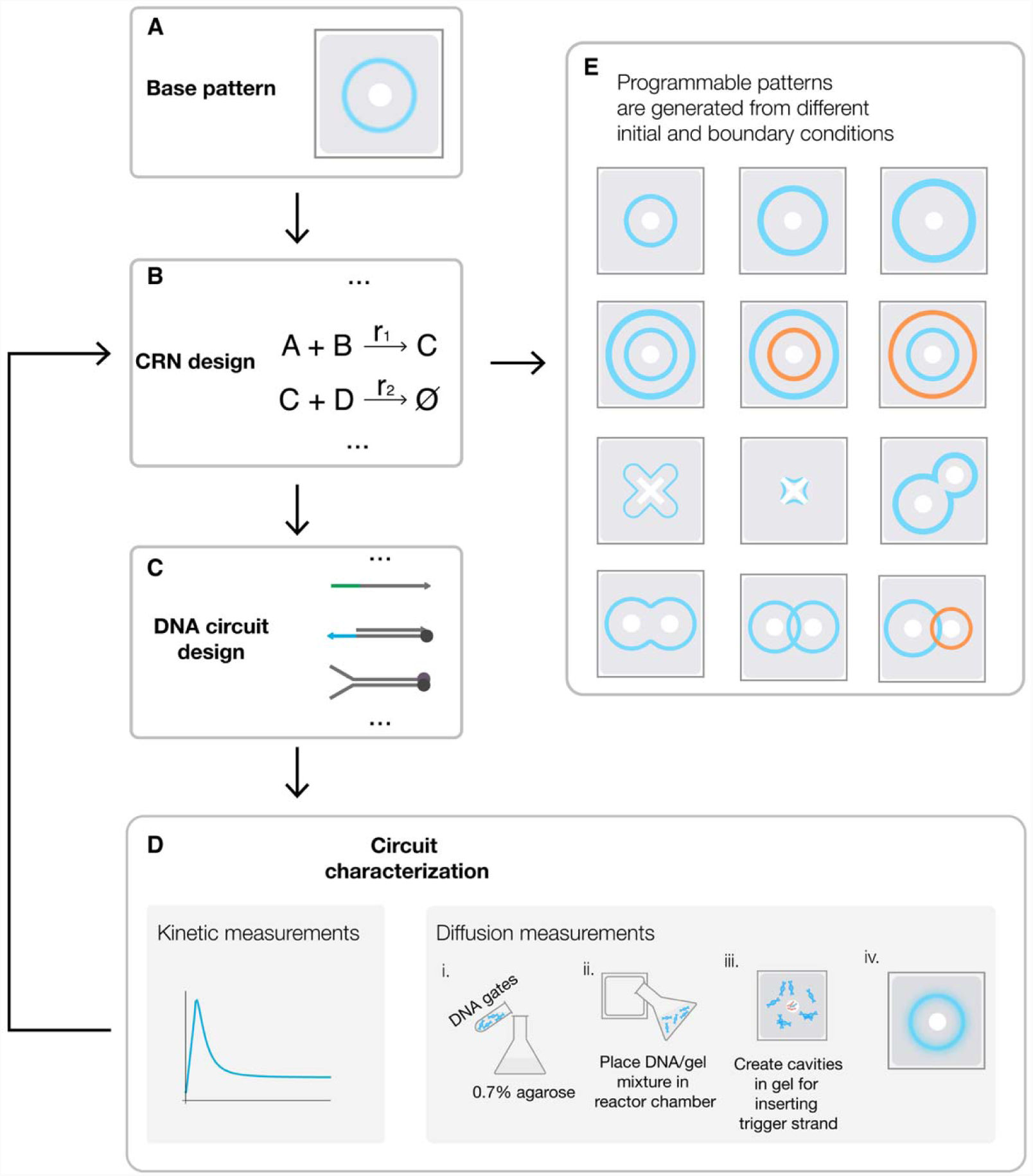
Overview of the workflow for DNA-based programmable patterning. **A.** We use a ring as the test case and basic building block for pattern formation. **B.** We designed a CRN to implement thi basic pattern. **C.** The CRN is compiled into molecules that realize the desired reaction. **D.** We performed experiments both in solution and in gel. Spectrometry experiments were used to measure parameters such as reaction constants and diffusion coefficients. These parameters were then entered as input to computer simulations for predicting the spatial dynamics of the system. The model also helps us to determine the initial conditions required for generating ringed patterns with target geometries. Gel measurements were conducted by suspending DNA gates in a hydrogel solution and molding the mixture into thin sheets in cast. To trigger programmed pattern formation, we loaded initiator strands into cavities in the DNA-hydrogel. Depending on the initial (concentrations of initiator strands) and boundary (shape and placement of cavities) conditions, gels embedded with identical DNA gates can be programmed to display different spatial dynamics. **E**. More complex patterns can be constructed by combining multiple ring-forming systems. The reaction diffusion model makes it possible to quantitatively simulate pattern formation before experimental implementation.

The core CRN for pulse formation consists of three reactants: activator, reporter, and inhibitor (**Fig. 2A**). An activator is a single-stranded DNA molecule used to initiate a reaction cascade. A reporter is a partially double-stranded DNA complex with a fluorophore-labeled signal (top) strand and a quencher-labeled bottom strand that is fully complementary to the activator. An inhibitor is a partially double-stranded DNA complex with an unmodified protector strand and a longer quencher strand that is fully complementary to the signal strand (**Fig. 2B**). We entered the desired domain structures into NUPACK Design to generate compatible sequences for building the pulse module (**Supplementary Section 1**). These sequences are listed in **Supplementary Table 1**. In a well-mixed setting, this three-component reaction module produces a single pulse via a two-step reaction (**Fig. 2B**): first, activators trigger fluorescence by releasing signal strands from reporters through toehold-meditated strand displacement; then, the signal is absorbed and repressed by the inhibitor.

**Figure 2.**
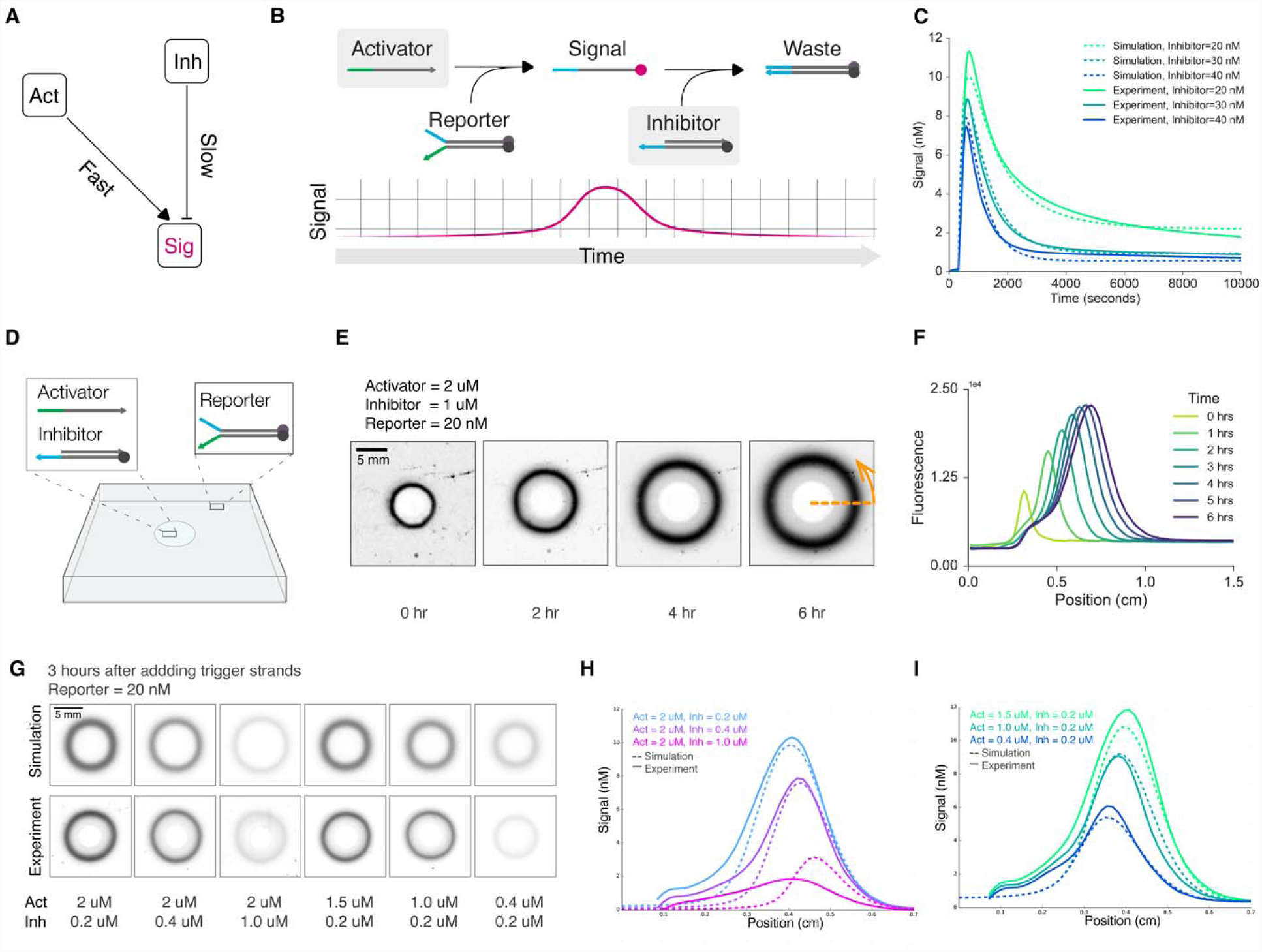
Single-ring pattern formation in a DNA-hydrogel. **A.** CRN for a pulse generator. Here, activators trigger the release of signals, while inhibitors repress signals. We designed the CRN such that signal activation always precedes signal inhibition. **B.** DNA strand displacement implementation of the CRN. Activators react with reporters to release signal strands. Fluorophores on the signal strands becom unquenched as they disengage from reporter complexes. Free floating signal strands can be absorbed by inhibitors, which suppress signals by quenching fluorophores. Because signal activation and inhibition occur sequentially, the observed fluorescent signal forms a pulse in time. **C.** Fluorospectrometry measurement of the DNA module (activator = 100 nM, reporter = 20 nM). Solid and dashed lines indicat experimental and fitted data, respectively. The duration and amplitude of the pulse can be tuned b varying the ratio of activators to inhibitors in the system. **D.** Configuration of the hydrogel experiment. A mixture of hydrogel and reporters was molded into a square sheet. A small circular cavity was made in the center of the sheet, where we loaded activators and inhibitors. **E.** Gel images showing a circular stripe pattern developing over the course of 6 hours. **F.** Intensity profiles of the same gel experiment. Th intensity profiles were obtained by taking the radially averaged intensity of gels at different time points. **G.** Varying the geometries of the single-ring pattern by changing the activator-to-inhibitor ratio. Top row: simulation results. Bottom row: gel experiment results. **H.** Varying inhibitor concentration while keeping activator concentration constant. **I.** Varying activator concentration while keeping inhibitor concentration constant.

To test the pulse module, we added reporter and inhibitor to a solution, triggered the reaction by adding the activator, and measured fluorescence changes using a spectrofluorometer. As designed, we can program the pulse shape by changing reactant concentrations. Specifically, pulse amplitude and duration depend on the rates of signal activation and inhibition: when we lowered the activator-to-inhibitor ratio in the solution, we observed a corresponding decrease in pulse amplitude and duration (**Fig. 2C**).

We developed a computational model in Visual DSD to simulate the pulse module (**Supplementary Sections 2 and 3**). The model consists of two reversible bimolecular reactions: signal activation (activator and reporter react to produce signal) and signal inhibition (signal is absorbed by inhibitor). Model parameters include reaction rate constants of signal activation and inhibition, as measured from separate spectrometry experiments (**Supplementary Figs. 1-4**). Our simulation confirms the module as a pulse generator (**Fig. 2C**) and further refines the rate constants to improve prediction accuracy (**Supplementary Section 3, Supplementary Table 2**).

## Programming single-ring patterns

Next, we set out to test spatial pattern formation. We synthesized DNA-hydrogel sheets by suspending reporters in 0.7% agarose solution. We used low melting point agarose and added reporter gates at room temperature to minimize denaturing. We cast the gels into thin sheets by pouring the mixed solutions into acrylic reactors. **Supplementary Figs. 5 and 6** illustrate this process in detail.

To initiate pattern formation, we loaded activators and inhibitors into a circular cavity at the center of the DNA-hydrogel sheet (**Fig. 2D**). As the diffusion fronts of activators and inhibitors advance, they react with the embedded reporters to trigger an outwardly propagating pulse, leading to the formation of a ring pattern. Thus, the ring pattern is a direct result of the interplay between diffusion and reaction. **Fig. 2E** shows the formation of a ring pattern over the course of 6 hours. Additionally, radially averaged intensity profiles provide quantitative information about pattern geometry not readily discernable from gel images alone. We found that the width and peak intensity of the ring grow over time as reporters are being triggered (**Fig. 2F**).

Like the amplitude and duration of a signal pulse measured in a well-mixed solution, the intensity and width of a ring also depend on the initial conditions of the DNA-hydrogel sheet. Reducing the initial concentration of activators resulted in rings with decreased widths and peak intensities (**Fig. 2G**). Radially averaged intensity profiles show that the inhibitor concentration controls the position of the trailing edge (**Fig. 2H**), while the activator concentration controls the position of the leading edge (**Fig. 2I**). To make meaningful comparisons, we established standard curves to convert fluorescence values to concentration units for spectrometry and gel image data (**Supplementary Figs. 7 and 8**).

Using Visual DSD, we built a predictive reaction-diffusion model to simulate pattern formation (**Supplementary Section 4**). The model uses rate constants inferred from spectrometry data (**Supplementary Table 2**) and assumes a common diffusion coefficient for all DNA complexes in our gel matrix (derivation of the diffusion coefficient is described in detail in **Supplementary Section 2** and **Supplementary Fig. 9**). The simulation results are displayed alongside corresponding gel images and show good quantitative agreement with the experimentally observed patterns (**Fig. 2G**).

## Building tunable concentric rings

Next, we asked whether we could control the radius of the ring pattern by adding a threshold component to our core CRN. The threshold is a single-stranded DNA that is fully complementary to the activator (**Fig. 3A**). Because hybridization between the activator and the threshold is faster than the reaction between the activator and the reporter, the threshold effectively acts as a sink to the activator.

**Figure 3.**
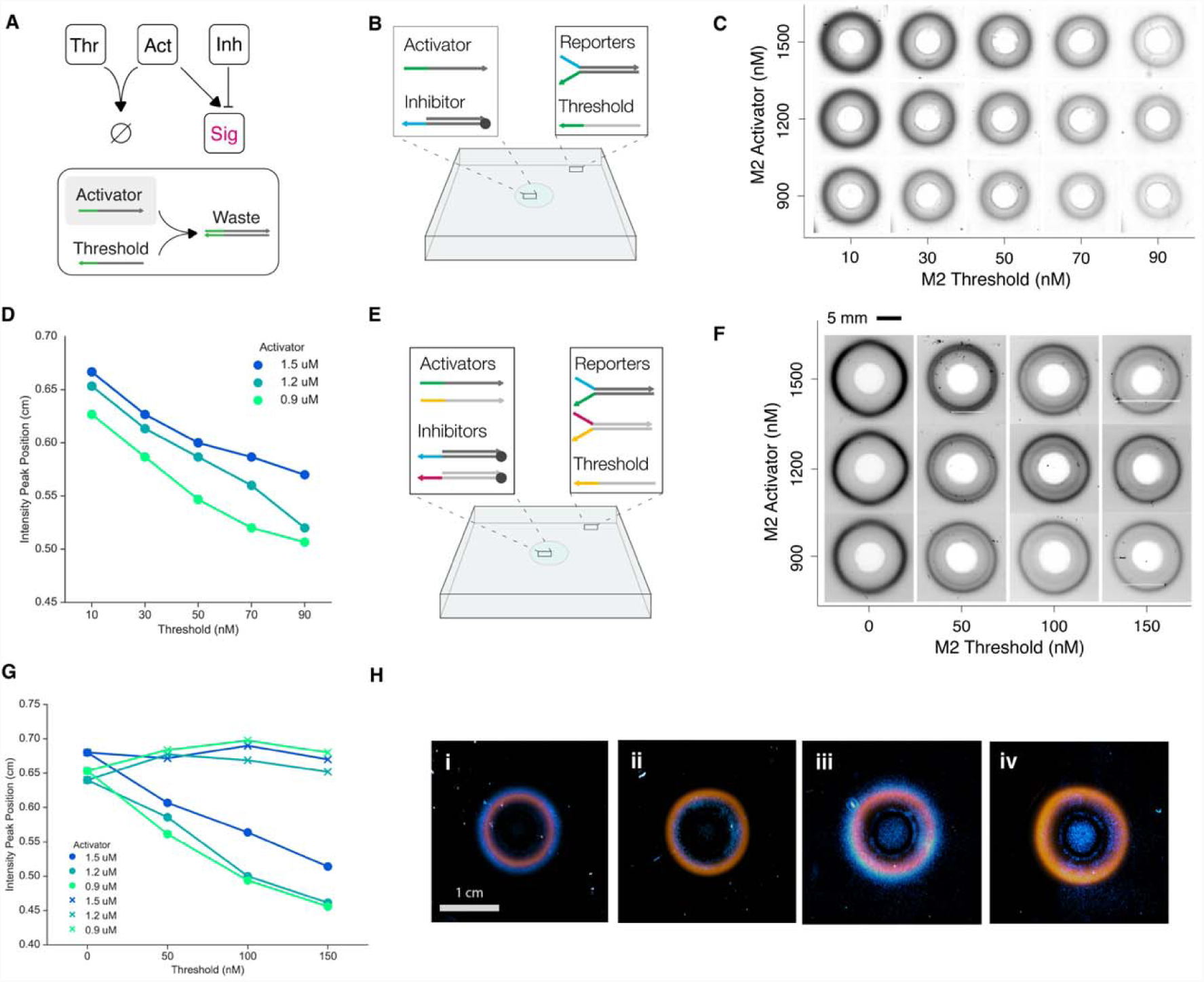
Tunable two-ring pattern formation. **A.** Schematics showing the pulse generating CRN and the corresponding DNA circuit modified to include a thresholding mechanism. Threshold strands act as sinks for activators to slow down the release of signals. **B.** Gel experiment setup for single-ring pattern with threshold. The hydrogel sheet is embedded with reporters and threshold. Activators and inhibitors are added to the circular cavity in the center of the sheet. **C.** Images showing gels embedded with five threshold concentrations at three activator concentrations. Inhibitor concentrations were set at 200 nM for every gel. Images were taken 3 hours after triggering. **D.** Intensity peak positions of threshold gel experiments. Increasing threshold concentration decreases ring radius. Changing the activator concentration has a visible, but much smaller effect on the radius. **E.** Gel experiment setup for concentric ring patterns. Each hydrogel is embedded with reporters from two orthogonal modules and threshold from only one module. To trigger the gel, we loaded the cavity with activators and inhibitors from both circuits. **F.** Images of concentric ring patterns in gels with two orthogonal modules. Each gel was synthesized according to the setup in **E**. We varied the threshold and activator concentrations for module M2 (reporter = 20 nM, inhibitor = 200 nM), while keeping concentrations for module M1 unchanged across gels (reporter = 20 nM, inhibitor = 200 nM, activator = 1200 nM). **G.** Intensity peak positions of concentric ring experiments. Peak positions for rings generated by module M1 (shown in “x” markers) remain stable across experiments, while peak positions for rings generated by module M2 (shown in circles) decrease linearly with increasing threshold. **H.** Programming the size and color of concentric ring patterns. We modified M2 such that its reporters are functionalized with Cy5 (in cyan) to differentiate it from M1 (FAM, in orange). We could selectively program the radius of a ring in a concentric ring pattern by controlling the concentration of threshold (gels 1 and 2). Changing the activator concentration has similar, but smaller, effect, so long as the gels contain non-zero concentrations of threshold (gels 3 and 4)

To study the effect of threshold in a spatial setting, we embedded threshold along with reporters in the hydrogel (**Fig. 3B**). Activators diffusing into the gel are annihilated upon encountering the threshold. Thus, signal activation only occurs once a region has been depleted of unreacted threshold. We prepared gels embedded with different threshold concentrations (**Fig. 3C, Supplementary Fig. 10**). We found that for gels triggered with identical activator and inhibitor concentrations, higher threshold concentrations reduced the radius proportionally; for gels embedded with a nonzero amount of threshold, decreasing the activator concentration also reduced ring radius, but less effectively than increasing the threshold **(Fig. 3D**). Here, we define the radius as the distance between the position of peak intensity and the center of the gel, as measured from radially averaged intensity profiles (**Supplementary Fig. 10**). We updated our predictive computational model to include the threshold, using empirically derived rate constants and diffusion coefficient (**Supplementary Fig. 11**). Comparisons between the radially averaged intensity profiles of gel images and spatial simulations show that the simulation performs well for predicting ring patterns under different initial conditions (**Supplementary Fig. 12**).

Leveraging our ability to program the ring radius, we proceeded to build programmable patterns of two concentric rings. We designed a second ring-forming module with the same components as the first module but orthogonal sequences (**Supplement Section 1, Supplementary Table 1**). Next, we combined the reporters for these two modules, labeled M1 and M2, in the same hydrogel and added threshold for M2 only (**Fig. 3E**). We prepared twelve DNA-hydrogels, corresponding to four M2 threshold and three M2 activator concentration levels (**Fig. 3F**), while maintaining the same concentrations of M1 components across all twelve gels. To trigger pattern formation, we loaded both M1 and M2 activators and inhibitors in the cavity. The outer ring’s radius remained largely unchanged across experiments. Meanwhile, the inner ring’s radius was proportional to the levels of activator and threshold, with high activator and low threshold values corresponding to larger radii (**Fig. 3G**). For gels without the threshold, we found that changing M2 activator concentrations alone had no effect on the radius of the inner ring (**Fig. 3G**), further validating the essential role of threshold for changing ring radius.

To better visualize the two-ring patterns, we replaced the FAM fluorophore in M2 with Cy5, such that M1 and M2 signals have distinct colors. We used this improved visualization to show that it is possible to program the order in which the rings appear. Using the same setup as the previous experiment, we prepared two gels containing either M1 threshold (**Fig. 3I**) or M2 threshold (**Fig. 3I**), but not both. The gels were subjected to identical conditions otherwise. In Gel I, M1 signal activation lags M2 signal activation, resulting in an orange ring (FAM, M1) encircled by a blue ring (Cy5, M2). In Gel II, the order of the rings is reversed. Alternatively, we embedded two gels with both M1 and M2 threshold and triggered them with either higher M1 activator (**Fig. 3I**) or higher M2 activator (**Fig. 3II**). This experiment shows that we can program the order of rings by either varying the concentration of initiator strands or the composition of the DNA hydrogel.

## Beyond isotropic patterns

So far, we have only considered programmable pattern formation with isotropic boundary conditions. Next, we went beyond this simple geometry and generated anisotropic patterns by changing the shapes and placements of cavities. First, we loaded activators into an “X” shaped cavity in a gel embedded with only reporters (**Fig. 4A**). Because reaction rates are proportional to the concentration of reactants, this gel configuration resulted in high signal at the center and low signal at the tips (**Fig. 4A**). We then amplified the asymmetry by embedding threshold in the hydrogels and, separately, by changing the angles between the legs of the X-shaped cavity. Adding threshold amplifies the time difference between signal activation at the tips and at the center, while decreasing the angle increases activator concentration in the interior of the angle, which leads to faster signal activation.

We prepared nine DNA-hydrogels divided into three groups based on their embedded threshold concentrations. Each group was further divided into three gels based on the angles in the “X” cavity on the gel (**Fig. 4B**). We measured the distance from the vertex to the center of the gel for different time points, cavity angles, and threshold concentrations (**Fig. 4C**). We also plotted the vertex distances for different time points and fitted the data to Fick’s equation to find the “effective diffusion coefficient”, a reaction-diffusion dependent parameter we use to quantify the speed of signal propagation (**Supplementary Fig. 15**). For the same angle, both vertex distance (at the last time point) and the effective diffusion coefficient show an approximately linear dependence in threshold concentration (**Fig. 4D**).

**Figure 4.**
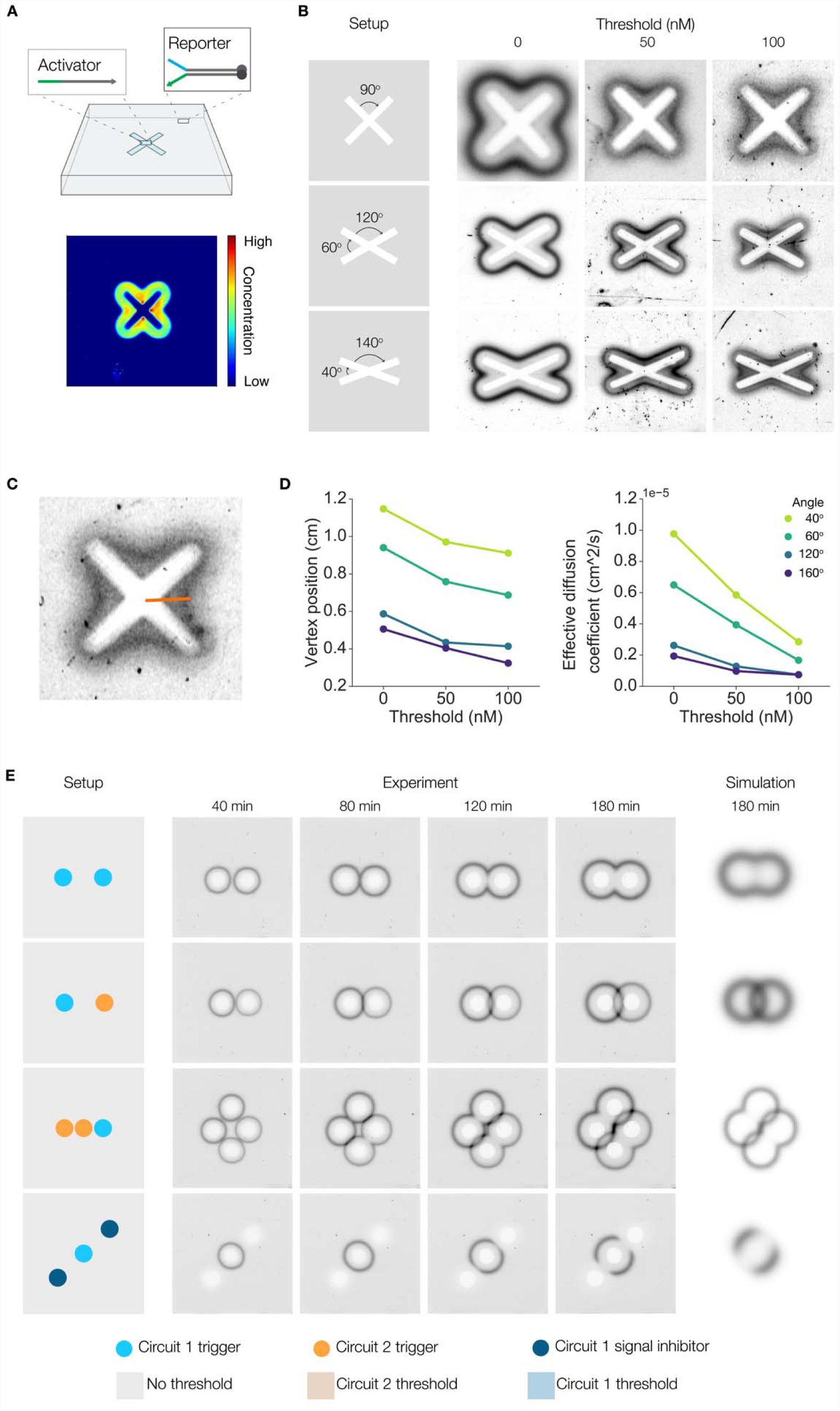
Programmed patterning from variations in boundary conditions. **A.** Asymmetrical cavities lead to non-isotropic concentration gradient in areas immediately surrounding the cavities. Points of greater curvature see higher concentrations of initiator strands. When activators were pipetted into an X-shaped cavity punctured on a gel containing only reporter gates (top), the resulting gel image (bottom) shows higher signal activation near the center than at the tips. **B.** The degree of asymmetry can be tuned by changing the level of threshold or the angle of curvature at the center. Higher threshold concentration results in a larger lag between signal activation times at the center than at the tips. Smaller angles of curvature lead to higher concentrations of initiator strands in the interior of the angle, which results in faster signal activation. **C.** We use the distance between the vertex and center of the gel as a proxy for how fast signal gets activated for each of the conditions tested. **D.** The diffusion coefficients were derived from measuring the distance between a signal vertex and the center at different time points (left). The dependence of the effective diffusion coefficients on threshold levels and angles of curvature is shown in the right plot. **E**. Patterns can also be programmed via the content and placement of cavities. We can induce conditional interference between signal fronts by changing the types and concentrations of initiator strands in the cavities. Gels I-IV were cast from the same DNA-hydrogel mixture yet evolved distinct patterns because they had different initial and boundary conditions. The DNA complexes loaded into each cavity are color coded and shown at the bottom of the figure.

Finally, we generated ring interference patterns by placing cavities at multiple locations in the gel. **Supplementary Table 3** lists the setup of each experiment in detail. Gels I-IV in **Fig. 4D** were synthesized from the same DNA-hydrogel solution containing M1 and M2 reporters. We applied different initial and boundary conditions to each gel to generate distinct patterns. For gel I, we loaded M1 activators and inhibitors in both cavities. We found that signals interfered destructively in regions where the rings intersected. This happens because the areas enclosed by the rings are devoid of reporters but are replete with inhibitors. Hence, as the two rings emanating from different cavities collide, their signal strands get consumed by the inhibitors from the opposite cavity. In contrast, when we loaded one of the cavities with M2 activators and inhibitors, we observed constructive interference of signals because the two diffusion fronts carry orthogonal reactants. Gel III was configured to display a combination of constructive and destructive signal interference. We created 3 cavities for gel IV. The center cavity was loaded with circuit 1 activators and inhibitors; the two peripheral cavities were loaded with circuit 1 inhibitors only. Since the signal and inhibitor diffusion fronts will annihilate each other wherever they collide, the resulting pattern is an incomplete ring with openings facing the directions of the peripheral cavities. **Supplementary Fig. 15** shows gels V and VI, where we used color coding and thresholding to induce two orthogonal rings of different radii that intersect.

## Discussion

We demonstrated a new approach to engineering programmable materials at the macroscale, using the reaction and diffusion of synthetic DNA strands to achieve quantitative and modular control over spatial patterns in hydrogels. To show proof of concept, we focused on a relatively simple pattern generated by non-catalytic CRNs. Incorporating more complex reaction networks, such as introducing feedback and cascading mechanisms, would produce more varied patterns^40^. We could also control diffusion by embedding appropriate complementary strands in the hydrogel to selectively slow down the diffusion of target DNA complexes, a possibility that has been explored in previous work^43^. Similar mechanisms could be used to convert transient patterns into permanent patterns by immobilizing signal strands with complementary capture sequences. Our system still relies on external spatial input and is therefore not fully autonomous and self-organizing. However, it is conceivable that a similar approach could be used to realize Turing patterns by combining nonlinear dynamics with control over diffusion rates.

Scalise *et al.* proposed making complex DNA-based programmable patterns by sequentially applying modular filters to an initially simple input pattern^41^. Our work has taken steps toward experimentally realizing such systems by predictably transforming simple input patterns into more complex output patterns. Thus, the workflow presented here can serve as an experimental basis for future projects exploring more complicated patterning systems.

In the longer term, we foresee applications where integrating chemical computing with additive manufacturing could expand the functionalities of existing 3D printed biomimetic materials. One could also imagine substituting fluorophores with other functional molecules, such as nanoparticles^44^ and quantum dots^45^, to synthesize novel materials with useful properties. Our work expands on previous research efforts in synthetic chemistry, synthetic biology, and DNA nanotechnology. Yet, we have only scratched the surface of the great array of programmable, macroscopic patterns and structures achievable by a synthetic DNA-based reaction-diffusion system. We believe this research presents a convincing case for using chemical computing in developing programmable matter.

## Acknowledgements

We would like to thank Neil Dalchau and Andrew Phillips from Microsoft Research for their valuable inputs on computational modeling. We also thank Yuan-Jyue Chen, Randolph Lopez, and Gourab Chatterjee for providing helpful advice. This work was support by ONR Award N000141612139 to G.S.

## Author Contributions

S.C. and G.S. conceived and developed the project. S.C. performed experiments and computer simulations. S.C. and G.S. analyzed the data and wrote the manuscript.

## Declaration of Interests

The authors declare no competing interests.

