## Supplementary Materials for "Programmable patterns in a DNA-based reaction-diffusion system"

Supplementary Figure 1: reaction kinetics for reporter activation	11

Supplementary Figure 2: reaction kinetics for signal inhibition.	11

Supplementary Figure 3: reaction kinetics of pulse response.	12

Supplementary Figure 4: reaction kinetics of adding threshold to reporter activation.	12

### S1 Module design

A single ring module consists of a maximum of four components: Activator, Inhibitor, Threshold, and Reporter. It is known that an incoherent feedforward loop can produce a single pulse. However, we decided to design a *de novo* module to minimize the number of components required, since fewer components and reactions reduce chances of crosstalk and simplify both the sequence design and simulation processes. Our guideline for module design is as follows. First, the module should have a minimal number of components and exhibit pulse behavior. Second, DNA complexes should have similar lengths such that a common diffusion coefficient could be used for all complexes. Third, each complex should have a minimal number of functional domains to simplify parameter inference and spatial simulation *in silico*. Following this guideline, we designed a pulse-generator containing three components (Activator, Reporter, Inhibitor) and three functional domains (two toehold and one migration domains).

We designed two orthogonal modules for implementing concentric ring patterns. These modules have identical domain level designs but orthogonal sequences. Both modules use FAM fluorophore (excitation = 495 nm, emission = 520 nm) for signal reporting. For patterns involving two different colors, we also ordered Module 2 reporters functionalized with Cy5 (excitation = 648 nm, emission = 668 nm).

We performed sequence selection using the design utility in NUPACK, which provided sequences based on the desired domain level designs. Subsequently, we entered these sequences into the analysis utility in NUPACK to validate that the sequences have minimal cross talk and obtained an estimate of rate constants based on the free energy differences in reactions ($\Delta\Delta G)$.

One concern we had was that the forked toehold in the reporter might lead to leak between the reporter and the inhibitor complexes. However, in solution experiment show that such leakage is essentially nonexistent.

### S2 Materials and methods

#### S1.1 DNA sequence design

All synthetic DNA gates used in this work have two functional domains: a single toehold domain and a single branch migration domain. We minimized the structural variations between gates to reduce simulation time and complexity. All toehold domains have 6 bp for optimal strand displacement speeds. All migration domains have 15 bp to optimize for sequence length and stability of the double strand complex.

#### S1.2 DNA gates preparation

Lyophilized synthetic DNA oligonucleotides purchased from IDT DNA were resuspended to 100 uM in 1X TAE Mg++ buffer and stored at -20^o^C. Single strands like the activator and the threshold were diluted to ~ 10 uM before use.

Concentrations of DNA were measured using a NanoDrop. For non-functionalized single strand DNA, we entered the sequence into the “oligo” mode in the NanoDrop software.

For partially double stranded complexes like the reporter and the inhibitor, we anneal their respective top and bottoms strands in 1:1 ratio at 20 uM in PCR tubes for a final volume of 100 uL. We anneal the strands in a thermocycler by first heating them to 95^o^C for 2 minutes, gradually cooling to 20^o^C at a speed of -1^o^C/min, and cooling for 10 minutes at 4 ^o^C before finally retrieving them from the thermocycler. We dilute the annealed reporters and inhibitors in 1X TAE Mg++ to 4 uM and 10 uM, respectively, and store at room temperature in the dark until use.

#### S1.3 Spectrometry measurement

Kinetics measurements were performed using a spectrofluorometer (Horiba FluoroMax4). The spectrometer can accommodate up to 4 cuvettes (0.875 ml Fluorometer Micro Square Cells).

FAM fluorophores have excitation and emissions wavelengths at 495 nm and 520 nm, respectively. The spectrometer was set accordingly to detect FAM functionalized oligos. We set the shutter width to 3 nm and the integration time to 10 seconds, with 5 seconds in between each cuvette measurement. Prior to each experiment, the cuvettes were cleaned three times with 70% ethanol followed by nine washes with deionized water, one final ethanol wash, and air dried.

For each cuvette, 1X TAE Mg++ buffer and reporter gates were mixed for a reporter concentration of 20 nM and a final volume of 700 uL. Before triggering the reporters, we ran the spectrometer for 15 minutes to establish a baseline for the background fluorescence. The spectrometer was paused to add activator and inhibitor gates at the requisite concentrations. An external water bath was connected to the spectrometer for the full duration of the experiment to keep the samples at a constant temperature of 25^o^C.

To normalize the spectrometry data, we measured the fluorescence of signal strands at concentrations of 10 nM, 20 nM, 30 nM, and 40 nM. We performed the same experiment on different days to ensure consistency and found minimal variation between measurements. We used this data to establish a standardized curve for converting fluorescence units to concentration (nM) and applied this conversion to all our spectrometry measurements.

#### S1.4 Making reaction chamber

We made a 3x3 grid on a large ¼’’ thick clear acrylic sheet by cutting out nine 4.5 cm x 4.5 cm squares on the sheet using a laser cutter. We chose these dimensions to ensure that the assembled reactor chamber could fit into the gel imager. To make the reactor chambers, we placed the grid over another ¼’’ thick clear acrylic sheet and taped the two sheets together on all four sides to create a leak-tight seal. We confirmed that the acrylic sheets are transparent when illuminated by light at the wavelengths of interest and did not interfere with the imaging of DNA-hydrogels.

To clean the reactor chambers after gel experiments, we first disassemble the grid and the underlying acrylic sheet by taking off the adhesives. After each gel experiment, we cleaned the grid and the sheet separately using liquid soap and warm water.

#### S1.5 Preparation of DNA-hydrogel sheets

To make 0.7% agarose gel, 0.35 g of low melting point agarose powder was added to 50 mL of 1X TAE Mg++ buffer in an Erlenmeyer flask and microwaved for 90 seconds or until the agarose had completely dissolved. Leftover hydrogel solution is stored at 37^o^C with plastic wrap covering the flask to minimize evaporation.

The gel solution is cooled at room temperature to below 37^o^C before synthetic DNA gates such as reporters and thresholds are added to prevent denaturing. After adding DNA complexes, we pipette 4 mL of DNA-hydrogel solution into a 5 mL Eppendorf tube and vortex on high for 10 seconds. Then the mixed solution is poured into a reaction chamber and chilled in a 4^o^C fridge for 15 minutes. Once gelation occurs, we move the gel back to room temperature. Next, we puncture circular cavities on the gel using the blunt end of a P10 pipette tip. We also made small 3D printed shapes as an alternative to using a pipette tip for creating cavities in the gel. The 3D printed parts include cylinders and X-shapes set at different intersecting angles. Each cavity can hold a volume of 40 uL.

#### S1.6 Gel imaging

We visualized the hydrogels using a Biorad PharosX gel imager. A 488 nm excitation laser and a 530 nm filter were used to image FAM fluorophores; a 637 nm excitation and a 695 nm filter were used to image Cy5 fluorophores. We scanned the images at 100 um resolution with the photomultiplier tube voltage at 80% of maximum. To make time course measurements, we installed a software (Mouse Recorder) that recorded specific mouse and keyboard actions to automatically take gel images at 30 minute intervals. Since gels dehydrate over time, we limited each gel experiment to less than 8 hours. When measuring gels containing both FAM and Cy5 fluorophores, we imaged the FAM channel first, then switched the filter and laser wavelength to the Cy5 channel.

#### S1.7 Gel image normalization

To create a calibration curve for converting gel imager fluorescence to concentration, we made six gels containing known concentrations of FAM functionalized signal strands (0 nM, 5 nM, 10 nM, 15 nM, 20 nM) and imaged these gels using the same imager settings as described in S1.6. We used ImageJ to calculate the average intensity in each gel, where we excluded the edges of the gels from intensity calculations because the gel is thicker there due to surface tension. We plotted the average fluorescence intensities against the known signal strand concentrations and fitted the data to obtain the equation for converting from fluorescence units to concentration (nM).

#### S1.8 Diffusion measurement

Because our oligos have the same length, we expect differences in diffusion rate to come from whether the DNA is single or double stranded. We used the diffusion coefficient of the signal strand as a proxy for other single strands in our system. For double strands, we use a modified reporter where the bottom strand is not functionalized with a quencher. We measured the diffusion coefficients of module M1 reporters and signal strand. For simplicity, we took the averaged value of the two diffusion coefficients and used it as the default value for all species in spatial modeling. To measure the diffusion coefficient, we pipetted 20 nM of either reporter or signal strand into a circular cavity punctured in a hydrogel made of 0.7% agarose only. The gel has the same dimensions as the other DNA-hydrogels. We took gel images periodically as the DNA strands diffuse over time.

Next, we used ImageJ to create a radial average intensity profile for each gel and normalize using the calibration curve from S1.7. For each intensity profile, we measured the position of the FWHM. Next, we plotted the positions against time and calculated fit to the plot using the 2D Einstein relation:

$$x^{2}=4Dt$$

, where *x* is the displacement (in our case, the position of the FWHM), *D* is the diffusion coefficient, and *t* is the time. The diffusion coefficients were later used in Visual DSD for spatial modeling.

The fitted diffusion coefficients are 5.6E-10 m^2^/s for the reporter complex and 6.6E-10 m^2^/s for the signal strand. These values are approximately five times greater than values for 20-bp DNA in free solution obtained from capillary electrophoretic measurements^1^. But because our strands diffuse in 2D agarose gels, the results here may not necessarily be comparable due to differences in experiment setup.

#### S1.9 Gel image analysis and processing

We analyzed the gel images (saved as TIFF files) on ImageJ. First, we performed auto-leveling on the images to increase contrast. This step affected the appearance of the images but not the intensity measurements. Next, we set the length scale such that measurements are in centimeters. An open source plug-in was used to carry out radially averaged intensity measurements.

For gels containing both FAM and Cy5 fluorophores, we took images taken at the same time point from both channels and combined them into a composite. Because the gel imager takes greyscale images by default, we colored the composite images in post-processing to enhance visual contrast.

### S3 Inferring reaction kinetics

#### S3.1 Chemical reaction network

We modeled the interactions between module components as the following set of bimolecular reactions:

$$Activator+Reporter \overset{\leftrightarrow}{k_{1r} , k_{1f}} Signal$$

$$Activator+Threshold \overset{\leftrightarrow}{k_{2r} , k_{2f}} Waste$$

$$Signal+Inhibitor \overset{\leftrightarrow}{k_{3r} , k_{3f}} Waste$$

#### S3.2 Strand displacement model

We constructed models of our DNA-based modules in Visual DSD, a compiler and programming language for describing DNA strand displacement reactions [cite]. The program is freely available at <https://dsd.azurewebsites.net/beta/>. We built the model to infer reaction rate constants using experimental data obtained from performing spectrometry on well-mixed solutions. First, we define the rate constants (k_1f_, k_1r_, k_2f_, k_2r_, k_3f_, k_3r_) and their range of possible values. For example:

k1f, (1.0e-7,2.0e-3)

Next, we define species in VDSD syntax, specifying the strands at the domain level:

def signal() = <t2^ d>	(*nM^-1^s^-1^*)

def activator() = <t1^ d>

def reporter() = <t2^>{d1^*}[d]

def inhibitor() = {t2^*}[d]

, where the angled, curly, and square brackets denote top-, bottom-, and double-strand complexes, respectively.

We also specified leak rates and concentrations of potentially faulty DNA complexes (due to annealing or stoichiometric errors) and their range of values (in units of nM):

good_i,(0.0,1.0)

good_a,(0.0,1.0)

bad_r,(0.0,10.0)

leak_i, (1E-07, 0.002)

We parallel the structure of the VDSD program with corresponding in solution experiments. For

experiments where we added in a DNA strand at a certain time point, we specified this condition with the following:

directive event activator() 1000.0*good_a @ T1

Here T1 denotes the time when activator was added. Parametrizing T1 is written as follows (units in seconds):

T1, (0.0,2000.0), 681.326827529902, real, random;

#### S3.3 Parameter inference

Using the model described in S2.1, we derived forward and reverse reaction rate constants for the four main reactions. We also fitted data for the full pulse-generator (reporter, inhibitor, activator, threshold). For the pulse-generator fits, we use rate constants derived from signal activation and inhibition spectrometry experiments as the initial values for parameter inference. The fit generates updated values of the activation and inhibition rate constants, which we use later in spatial simulations.

For each module, we tested all 5 reactions. We averaged the same reaction constant from different experiments. The result is used in simulating 2D spatial experiments.

### S4 Reaction-diffusion spatial simulations

To simulate 2D patterns, we built spatial models using the CRN mode in Visual DSD. Compared to the DSD mode, the CRN mode abstracts the interactions between module components as chemical reactions equations, without detailing the compositions of the reactants. We chose this simplified approach because it vastly reduces simulation time without compromising on simulation accuracy.

To construct the model, we specify the initial and boundary conditions using the syntax of Visual DSD. First, we define reaction rate constants using values inferred from spectrometry data (See Supplementary Table 3), where rate constants are expressed:

directive parameters [k1f = 1e-4; k1r = 1e-5; k2f = 5e-5; k2r = 1e-6]	(*nM^-1^s^-1^*)

Next, we define the spatiotemporal mesh and step sizes:

directive nx 101	(*step size, unitless*)

directive xmax 0.045	(*in meters*)

We define a common diffusion coefficient (2E-9 m^2^/s) for all strands since differences between strand sizes are negligible:

directive defaultdiffusion 6e-10

We use the following code to define the positions of cavities and the concentrations of initiating strands contained in the cavities:

directive spatialic centralcore {width=0.1; species = Act1; inner=2000.0}

Here, the first term in the curly bracket specifies the size of the cavity, defined in units of … The second term specifies the reactant in the cavity. The third term specifies the concentration of the reactant in units of nM.

Next, we describe the reactions and specify the appropriate rate constant:

Act1 + Rep1 ->{r2} S1 + W1 |

Inh1 + S1 ->{r2} W2 |

The two equations above describe the activation and inhibition of signals, respectively. Finally, we list the molecule type and their initial concentrations in the gel:

init Act1 2000 |

init Inh1 N*600 |

init Rep1

### Supplementary Figures

$$Activator+Reporter \overset{\leftrightarrow}{k_{1r} , k_{1f}} Signal$$

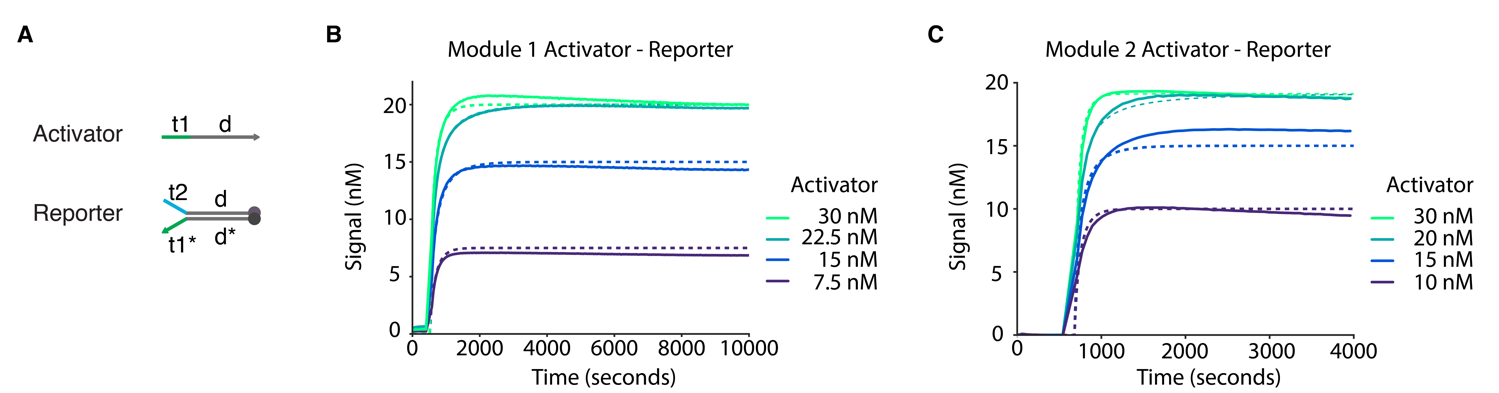

Supplementary Figure 1: reaction kinetics for reporter activation**.** We added 20 nM of reporter first to establish a baseline of background florescence. Then we paused the spectrometer to add to the indicated concentrations of activators. Solid lines show experimental data; dashed lines show simulated fits.

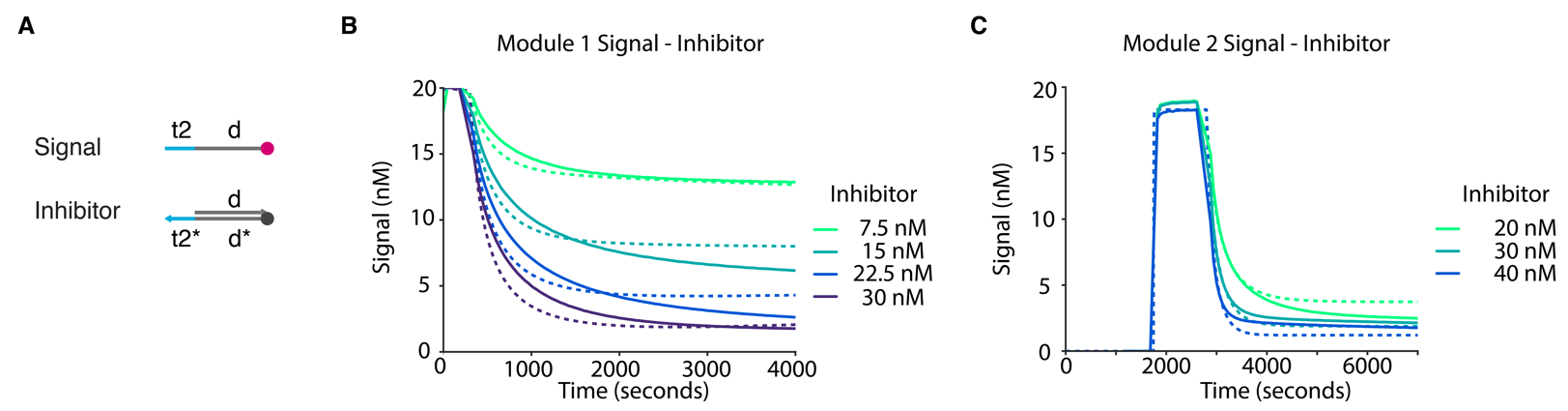

Supplementary Figure 2: reaction kinetics for signal inhibition. 20 nM of reporter was added to the buffer solution first, which was activated by 40 nM of activator. Finally, we added inhibitor at the indicated concentrations.

$$Signal+Inhibitor \overset{\leftrightarrow}{k_{3r} , k_{3f}} Waste$$

$$Activator+Reporter \overset{\leftrightarrow}{k_{1r} , k_{1f}} Signal$$

$$Signal+Inhibitor \overset{\leftrightarrow}{k_{3r} , k_{3f}} Waste$$

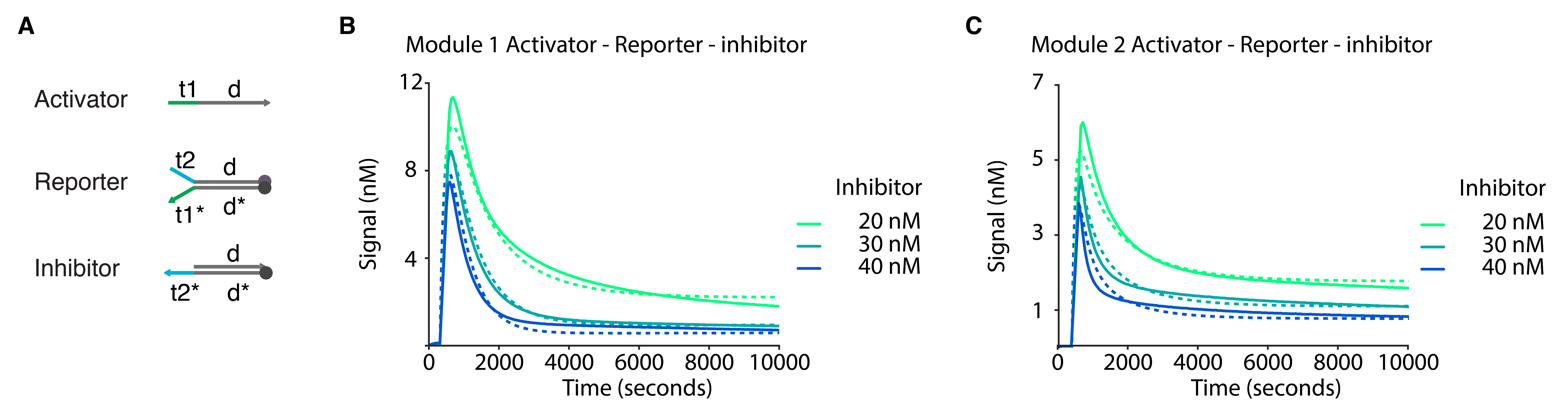

Supplementary Figure 3: reaction kinetics of pulse response. 40 nM of activator was added to each sample solution containing 20 nM of reporter and inhibitor at the indicated level of concentration.

$$Activator+Threshold \overset{\leftrightarrow}{k_{2r} , k_{2f}} Waste$$

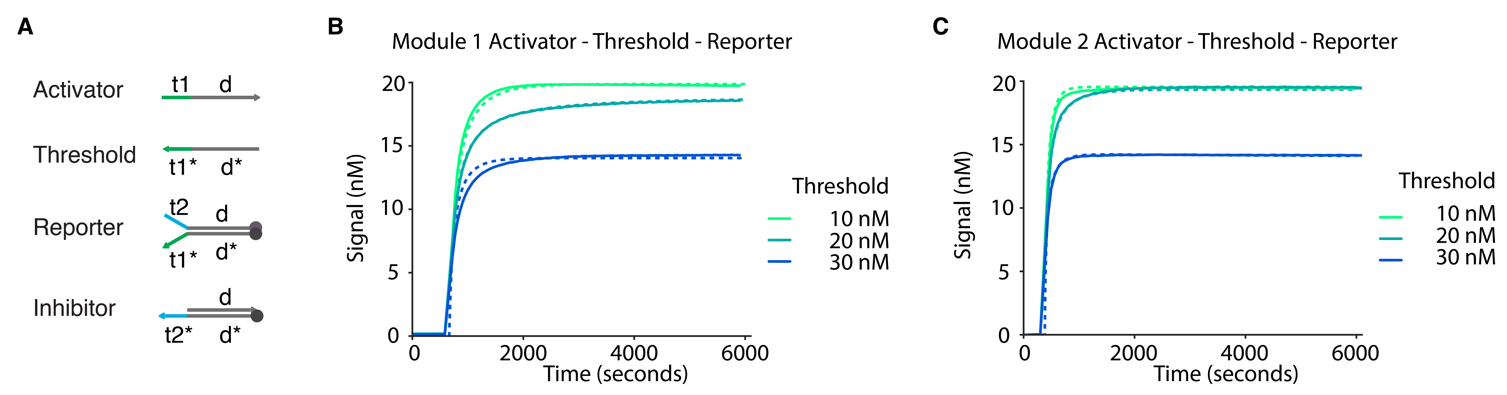

$$Activator+Reporter \overset{\leftrightarrow}{k_{1r} , k_{1f}} Signal$$

Supplementary Figure 4: reaction kinetics of adding threshold to reporter activation. 40 nM of activators were added to each sample solution containing 20 nM of reporter and threshold at the indicated concentrations.

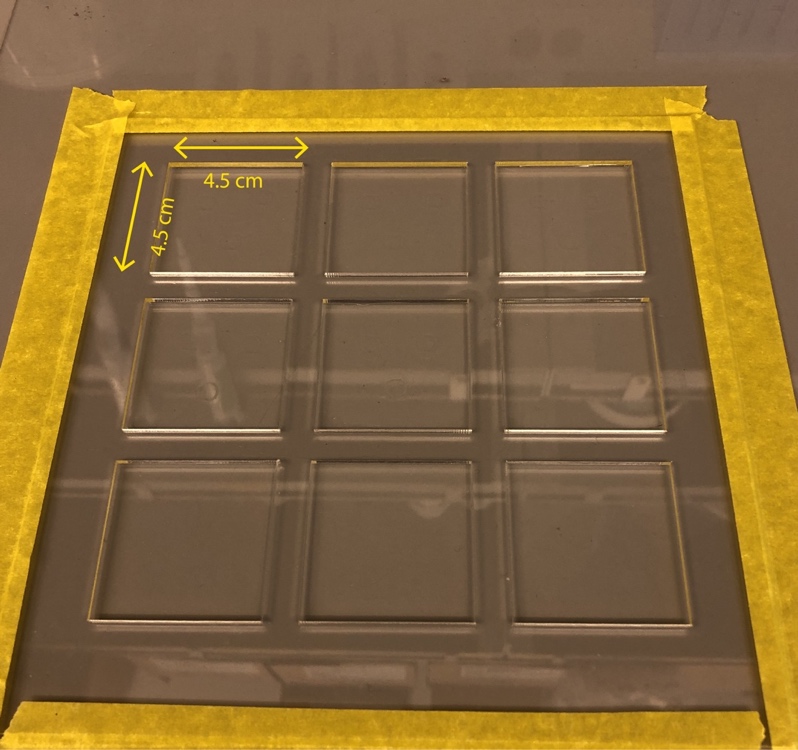

Supplementary Figure 5: acrylic reactor chamber setup. To make the reaction chamber, we taped two customized acrylic pieces together. The top piece contains nine grids measuring 4.5 cm x 4.5 cm. The bottom piece is a flat sheet of acrylic. Both sheets measure 0.25’’ in thickness.

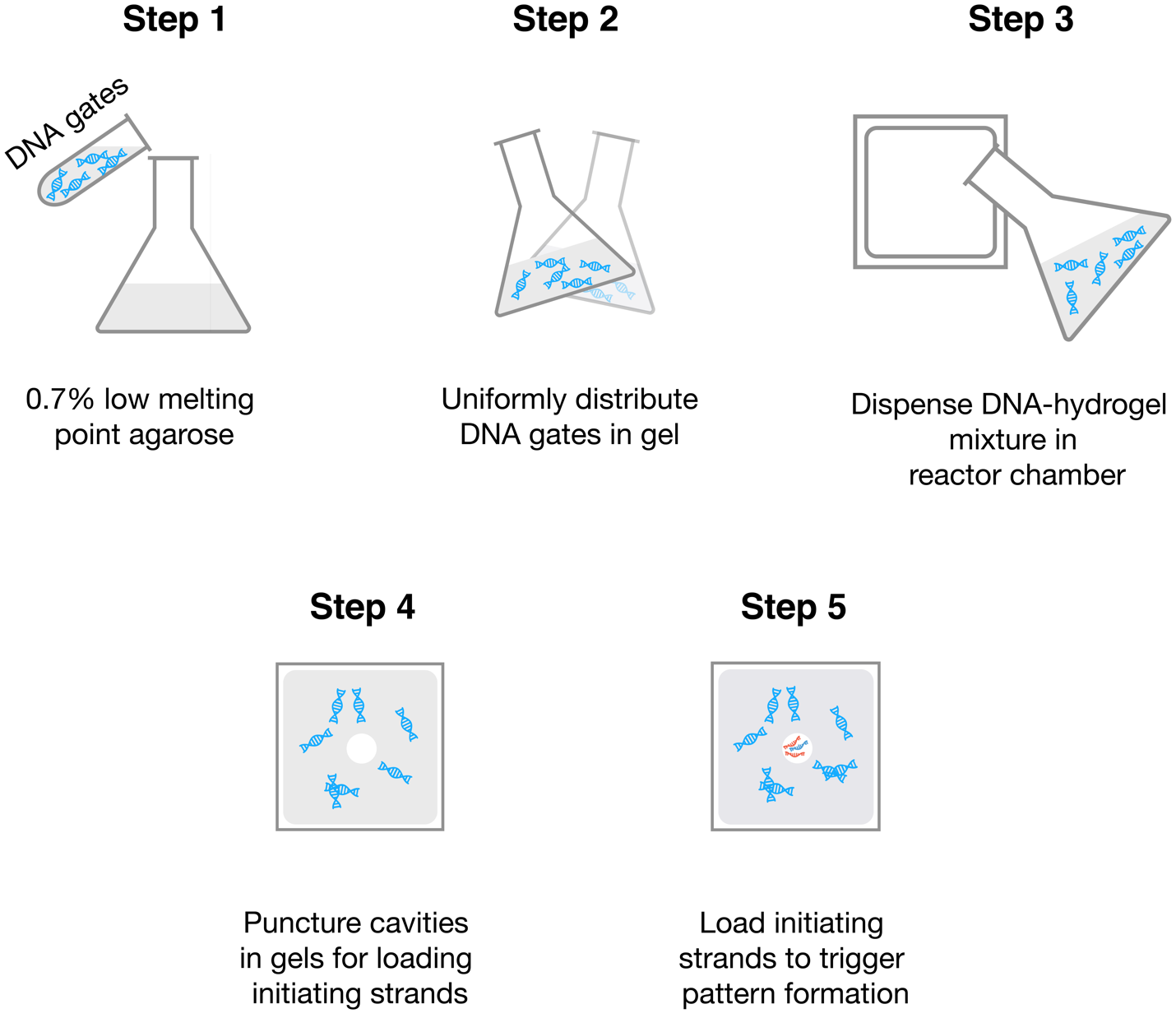

Supplementary Figure 6: workflow for synthesizing DNA-hydrogels. We used low melting point agarose to minimize denaturing of DNA complexes. After thoroughly mixing DNA with agarose gel solution, we dispense the mixture in acrylic reactor chambers. To make non-circular cavities, we placed 3D printed shapes in the hydrogel solution before gelation. To make circular cavities, we punctured directly on the hydrogel (post-gelation) using the blunt end of a 10 uL pipette tip. We pipetted initiator strands like activators and inhibitors into the cavities to trigger pattern formation. Each cavity can hold a maximum of 40 uL.

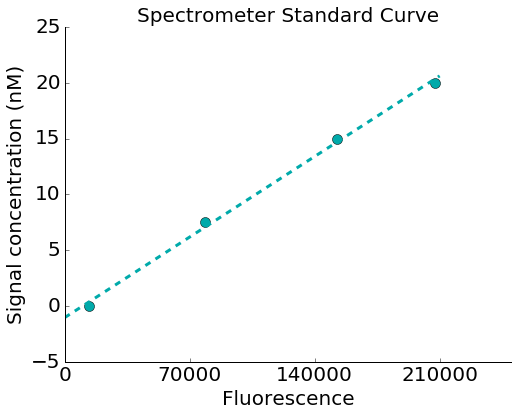

Supplementary Figure 7: calibration curve for spectrometer. Standard curve derived from measuring signal strand at four concentrations in a well-mixed solution. We convert fluorescence units to nM before performing inference on the data.

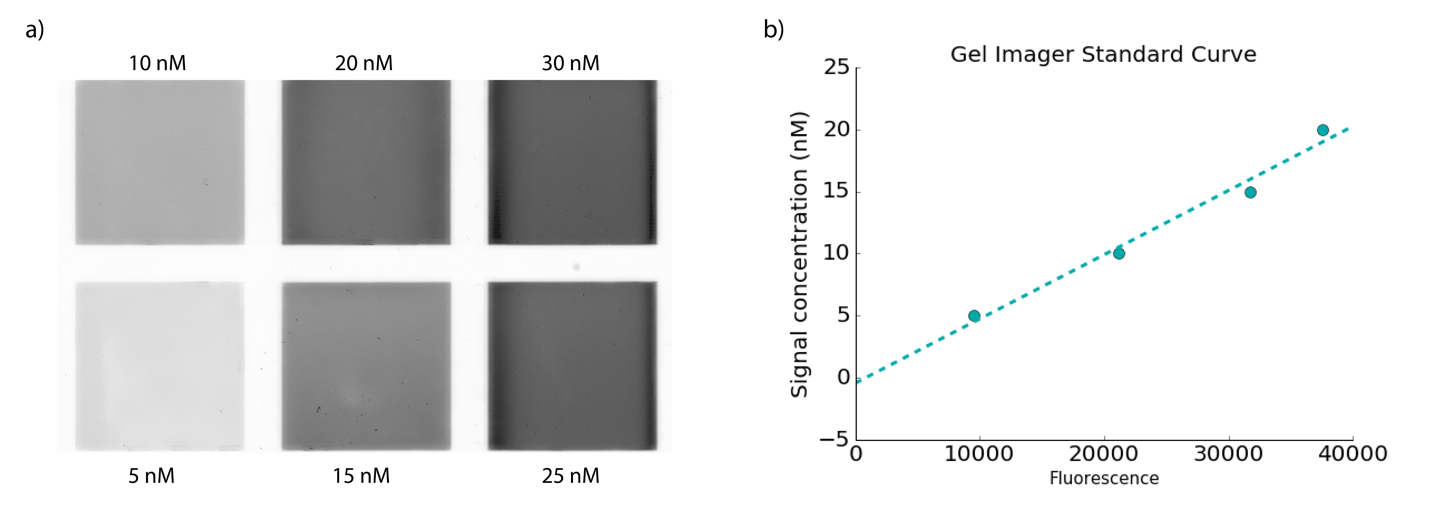

Supplementary Figure 8: calibration curve for gel imager. a) Gels uniformly embedded with the labeled concentrations of signal strands. b) Calibration curve for converting gel imager fluorescence intensity to signal strand concentrations.

|  | D (cm^2/hr) | D (m^2/s) |
| --- | --- | --- |
| Module 1 reporter complex | 0.020 | 5.6E-10 |
| Module 1 signal strand | 0.024 | 6.6E-10 |

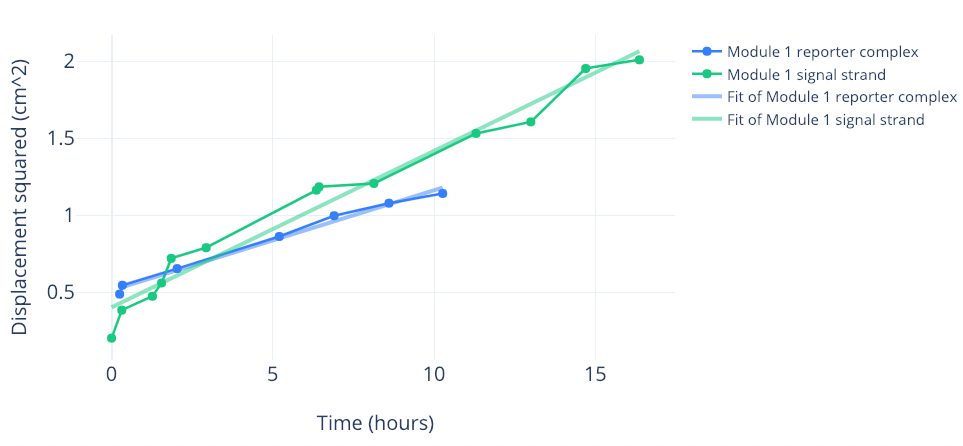

Supplementary Figure 9: diffusion coefficients of signal strand and reporter. For obtain each data point on the plot, we first measured the position of the FWHM on a radially averaged intensity profile at the specified time point, then calculated the square of this position. We plotted displacement squared against time and fitted the data to the 2D mean square displacement equation. The table at the top shows the diffusion coefficients for the signal strand and the report complex in units of cm^2^/hr and m^2^/s.

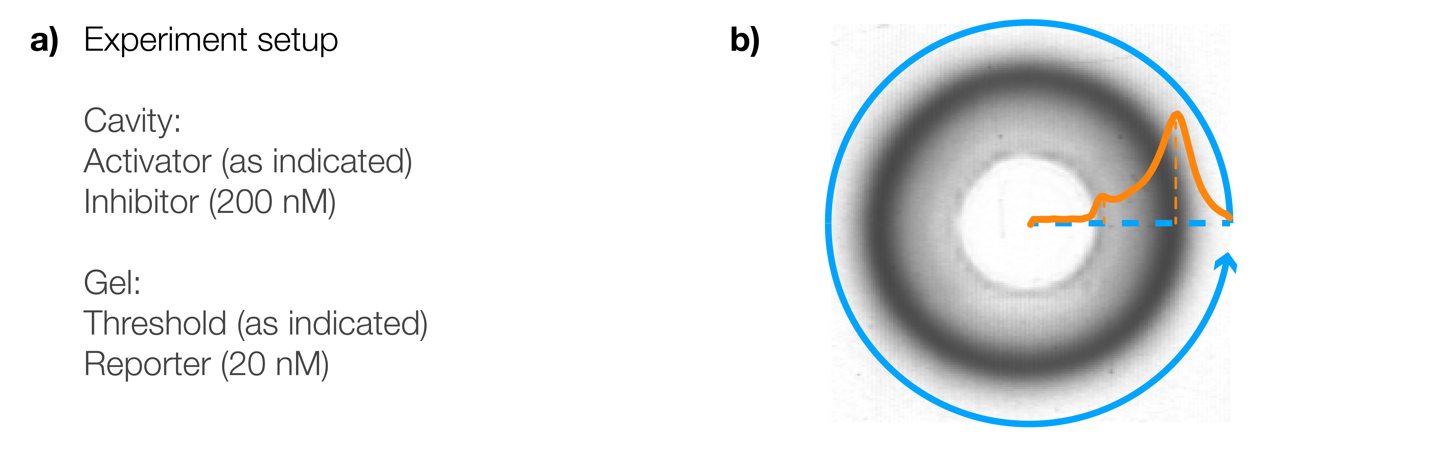

Supplementary Figure 10: experiment setup of single ring patterns in threshold-added gels. **a)** The concentrations of reactants and where they were added. Values of activator and threshold concentrations can be found in **Supplementary Fig. 12**. **b)** An example showing how we measured the radially averaged intensity profile using gel images. Two visible “peaks” are visible in each profile. The smaller peak corresponds to the edge of the cavity. The larger peak corresponds to signal pulse.

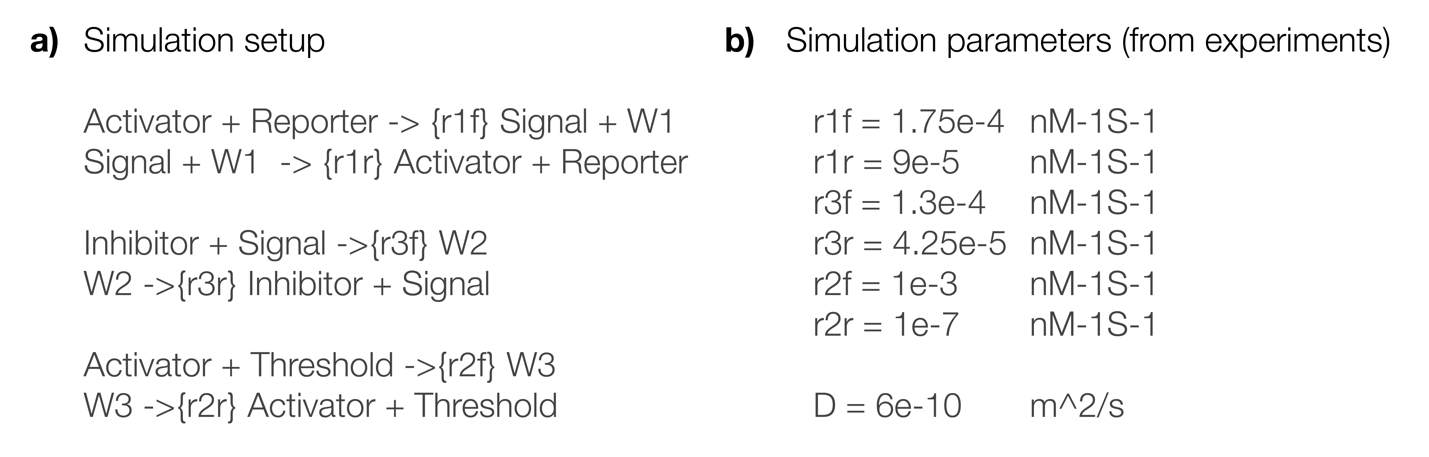

Supplementary Figure 11: simulation setup for single ring pattern in threshold-added gels. **a)** The chemical reactions for performing spatial simulations of single ring patterns in Visual DSD. The simulation includes the forward and reverse rate constants. **b)** The rate constants and diffusion coefficient used in the simulation. We derived these parameters from experiments.

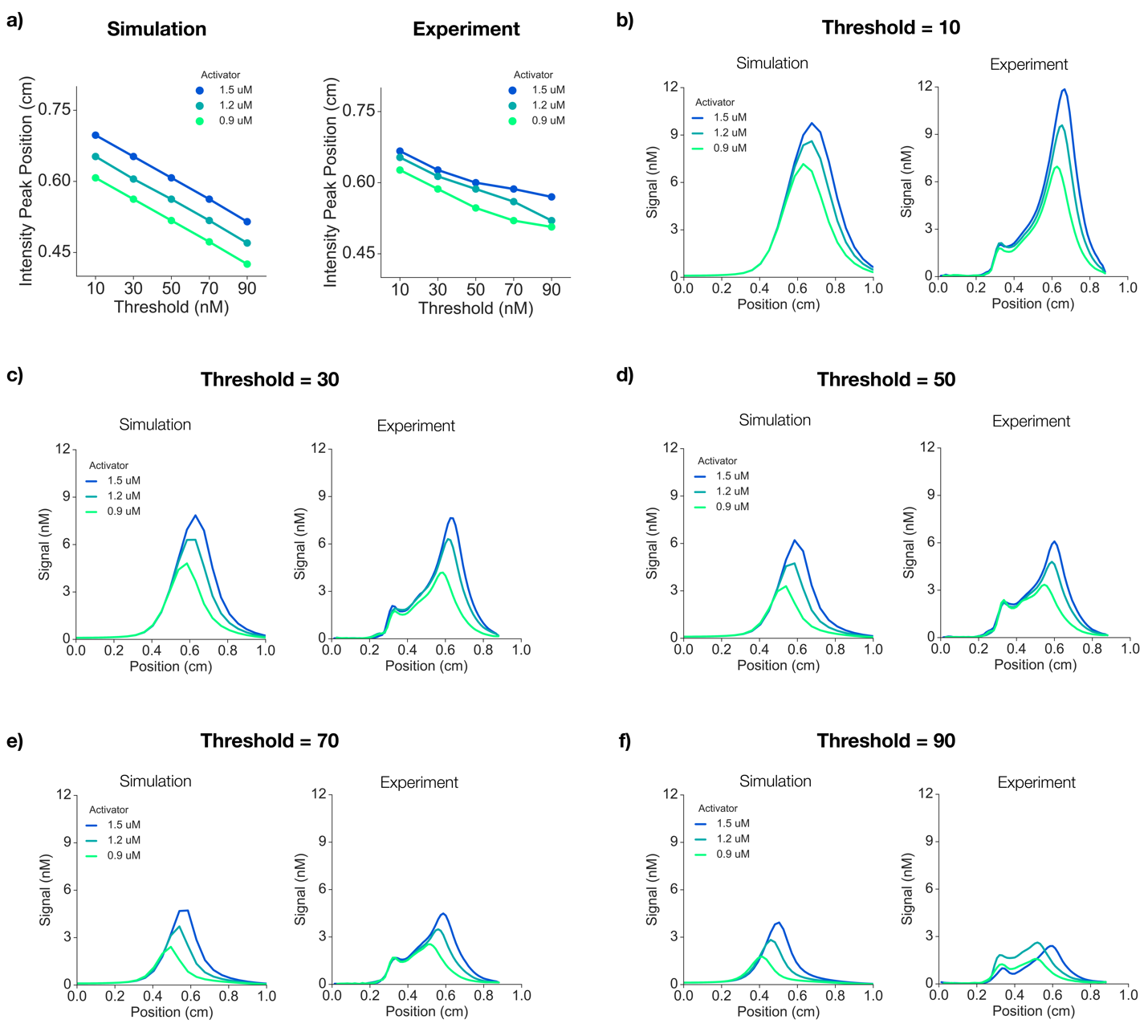

Supplementary Figure 12: single ring patterns in threshold-added gels (*in vitro* vs *in silico*). **a)** The threshold is more effective than the activator for changing the positions of intensity peaks, a proxy for ring radius. Each point on the plot represents the peak intensity position of a ring pattern, as measured from the corresponding radially averaged intensity profiles. **b)-f)** Comparisons between the radially averaged intensity profiles of simulated and experimentally realized ring patterns for the setups described in **Supplementary Figs. 10** and **11**.

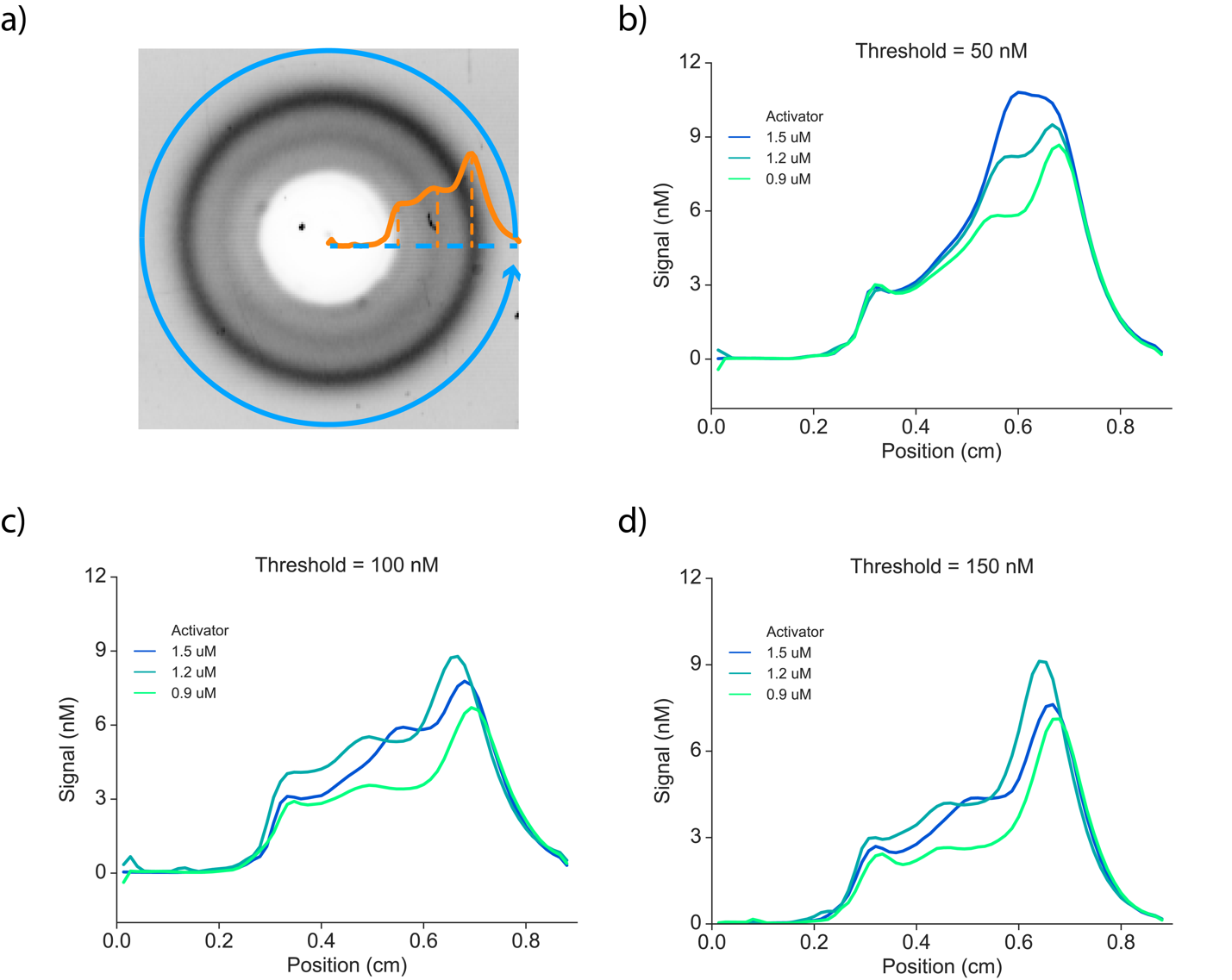

Supplementary Figure 13: intensity profiles for concentric ring patterns. **a)** Three visible “peaks” are visible in each profile. The smallest peak corresponds to the edge of the cavity. A second peak indicates the inner ring. The largest peak corresponds to the outer ring. **b)-d)** Each plot shows the intensity profiles from three gels embedded with the indicated concentration of threshold, but triggered with three different activator concentrations. For the inner ring intensity peaks, we measured the position where the signal showed the greatest inflection. The positions of intensity peaks and their dependence on threshold and activator

concentrations are shown in **Fig. 3G**.

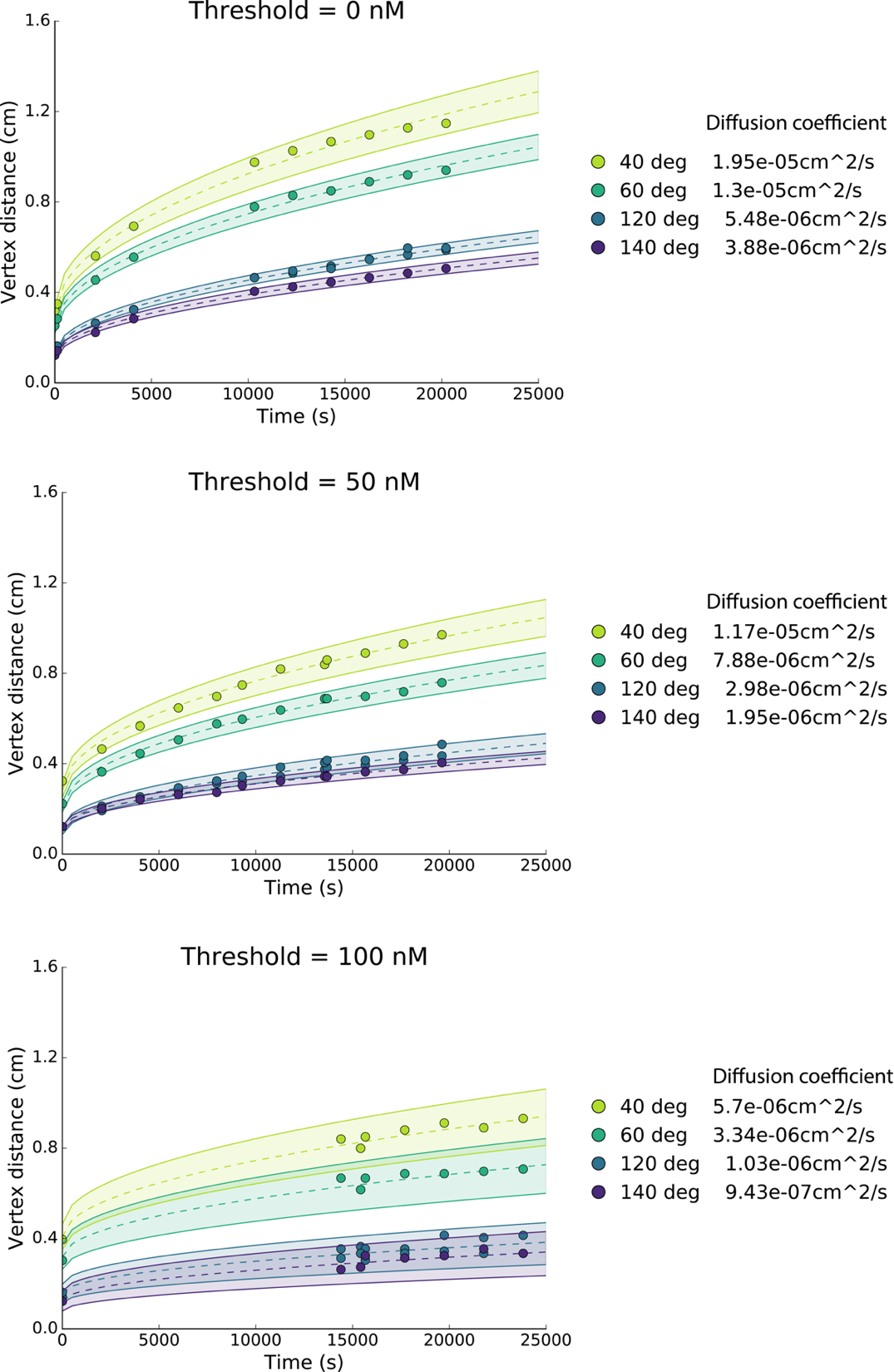
Supplementary Figure 14: angular and threshold on effective diffusion. The effective diffusion coefficient incorporates the reaction and diffusion aspects of the propagating signal. Each data point corresponds to the position of peak intensity relative to the center of the cavity. We fitted each dataset to the 2D Einstein equation to obtain the effective diffusion coefficient. For the last plot, the missing time points were due to malfunctions in the gel imager.

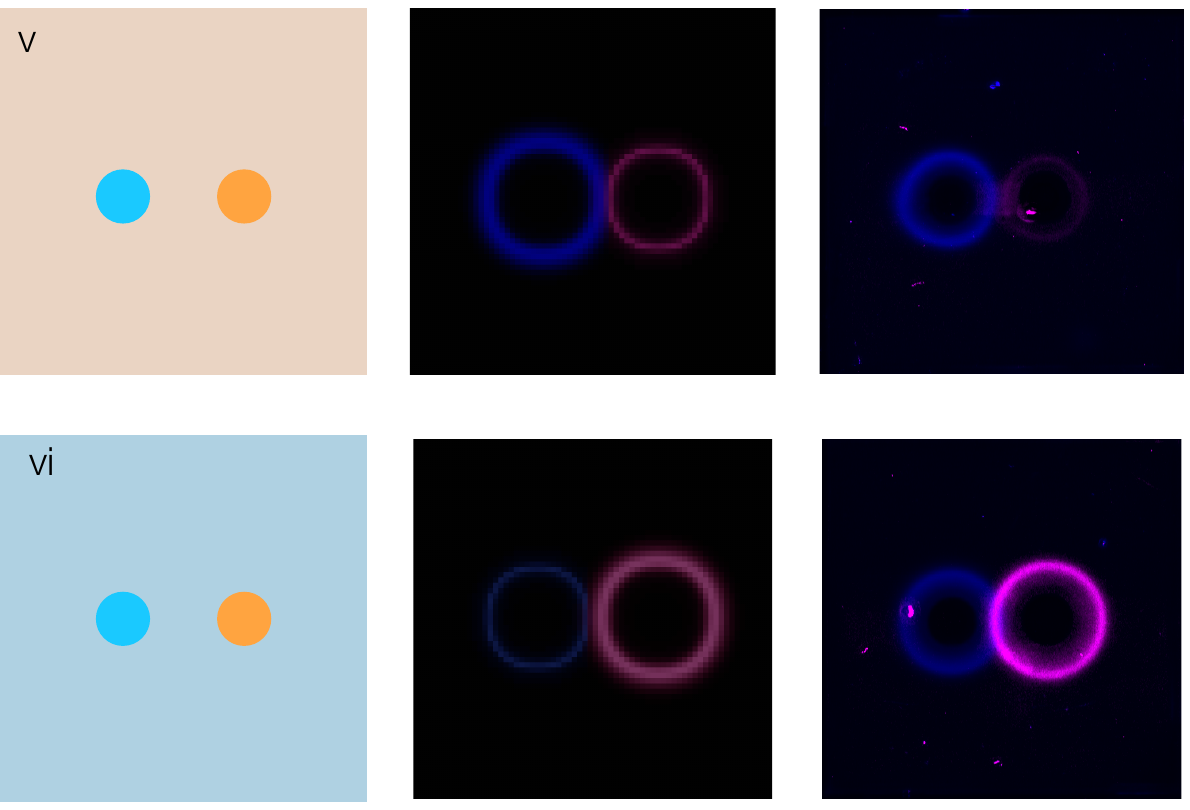

Supplementary Figure 15: two colored rings subjected to different threshold levels. Each gel was embedded with components from modules M1 and M2, which are orthogonal in sequence and have different reporter fluorophores. M1 reporters have FAM fluorophores (in magenta); M2 reporters have Cy5 fluorophores (in blue). The left column shows the gel setup; the center column shows simulation results; the right column shows gel image results. The top row and bottom rows show gels embedded with M1 and M2 threshold, respectively. For each gel, we pipetted M1 initiators in the left cavity and M2 initiators in the right cavity (activator = 2000 nM, inhibitor = 400 nM).

### Supplementary Tables

**Supplementary Table 1**

| Strand name | Sequence |
| --- | --- |
| AITR4.A | AAC CCA CAA AAC AAA ACC TCC |
| AITR4.T | GGA GGT TTT GTT TTG TGG GTT |
| AITR4.Rtop | CTT ATA CAA AAC AAA ACC TCC /36-FAM/ |
| AITR4.Rbot | /5IABkFQ/GGA GGT TTT GTT TTG TGG GTT |
| AITR4.Itop | CAA AAC AAA ACC TCC |
| AITR4.Ibot | /5IABkFQ/GGA GGT TTT GTT TTG TAT AAG |
| AITR5.A | CTT CTC CAT TCC TAC ATT TCC |
| AITR5.T | GGA AAT GTA GGA ATG GAG AAG |
| AITR5.Rtop.FAM | CAA TAT CAT TCC TAC ATT TCC/36-FAM/ |
| AITR5.Rbot.FAM | /5IABkFQ/GGA AAT GTA GGA ATG GAG AAG |
| AITR5.Rtop.Cy5 | CAA TAT CAT TCC TAC ATT TCC /3Cy5Sp/ |
| AITR5.Itop. | CAT TCC TAC ATT TCC |
| AITR5.Ibot.FAM | /5IABkFQ/GGA AAT GTA GGA ATG ATA TTG |
| AITR5.Ibot.Cy5 | /5IAbRQ/GGA AAT GTA GGA ATG ATA TTG |

**Supplementary Table 2:** rate constants obtained by fitting Visual DSD strand displacement model to spectrometry data. For rate constants that appear in multiple reactions, we provide the averaged value and the standard deviation.

| Rate constant (Module 1) | signal activation | Signal inhibition | signal activation and inhibition | signal activation with threshold | Average | Stdev |
| --- | --- | --- | --- | --- | --- | --- |
| k1f | 3.65E-04 |  | 1.61E-04 |  | 1.75E-04 | 1.83E-04 |
| k1r | 1.96E-04 |  | 4.90E-05 |  | 9.38E-05 | 8.88E-05 |
| k3f |  | 1.91E-04 | 7.13E-05 |  | 1.31E-04 |  |
| k3r |  | 2.51E-05 | 5.99E-05 |  | 4.25E-05 |  |
| Rate constant (Module 2) | signal activation | signal inhibition | signal activation with threshold | signal activation and inhibition | Average | Stdev |
| k1f | 9.63E-04 |  | 7.03E-04 | 1.35E-04 | 6.00E-04 | 4.23E-04 |
| k1r | 2.02E-04 |  | 1.72E-06 | 4.02E-04 | 2.02E-04 | 2.00E-04 |
| k2f |  |  | 1.96E-03 |  | 1.96E-03 |  |
| k2r |  |  | 1.06E-07 |  | 1.06E-07 |  |
| k3f |  | 1.92E-04 |  | 1.32E-04 | 1.62E-04 | 4.24E-05 |
| k3r |  | 1.68E-05 |  | 1.23E-05 | 1.46E-05 | 3.18E-06 |

**Supplementary Table 3:** detailed experiment setups for DNA-hydrogels

| Single ring experiment (Figure 2G) | |
| --- | --- |
| Gel | 20 nM reporter (M1) |
| Cavity | Activator (M1); Inhibitor (M1) |

| Threshold experiment (Figure 3C) | |
| --- | --- |
| Gel | 20 nM reporter (M2); Threshold (M2) |
| Cavity | Activator (M2); Inhibitor (M2) |

| Two concentric ring experiment (Figure 3F) | |
| --- | --- |
| Gel | 20 nM reporter (M1); 20 nM reporter (M2); Threshold (M2) |
| Cavity | Activator (M1); Inhibitor (M1); Activator (M2); Inhibitor (M2) |

| Two color experiments (Figure 3H, i&ii) | |
| --- | --- |
| Gel | 20 nM reporter (M1); 20 nM reporter (Cy5, M2); Threshold (M2) |
| Cavity | Activator (M1); Inhibitor (M1); Activator (M2); Inhibitor (M2) |

| Two color experiments (Figure 3H, iii&iv) | |
| --- | --- |
| Gel | 20 nM reporter (M1); 20 nM reporter (Cy5, M2); Threshold (M2) |
| Cavity | Activator (M1); Inhibitor (M1); Activator (M2); Inhibitor (M2) |

| Non-isotropic cavity experiments | |
| --- | --- |
| Gel | 20 nM reporter (M1) |
| Cavity | Activator (M1) |

| Non-isotropic cavity experiments | |
| --- | --- |
| Gel | 20 nM reporter (M1); Threshold (M1) |
| Cavity | Activator (M1) |

| Interference pattern experiments | |
| --- | --- |
| Gel | 20 nM reporter (M1); Threshold (M1) |
| Cavity | Activator (M1) |

### Supplementary Videos

**Supplementary video 1** Time course of single ring pattern for different activator and inhibitor concentration ratios as acquired over 5 hours (Fig. 2G).

**Supplementary video 2** Time course of concentric ring patterns for different activator concentrations with embedded threshold concentrations of 50 nM (Fig. 3F).

**Supplementary video 3** Time course of concentric ring patterns for different activator concentrations with embedded threshold concentrations of 100 nM (Fig. 3F).

**Supplementary video 4** Evolution of patterns corresponding to Fig. 4D first row.

**Supplementary video 5** Evolution of patterns corresponding Fig. 4D second row.

**Supplementary video 6** Evolution of patterns corresponding to Fig. 4D third row.

**Supplementary video 7** Evolution of patterns corresponding to Fig. 4D fourth row.

**Supplementary video 8** Evolution of patterns corresponding to Supplementary Fig. 15 (gel V).

**Supplementary video 9** Evolution of patterns corresponding to Supplementary Fig. 15 (gel VI).
