## Supplementary figures and images for "Programmable patterns in a DNA-based reaction-diffusion system"

### Supplementary Video 1

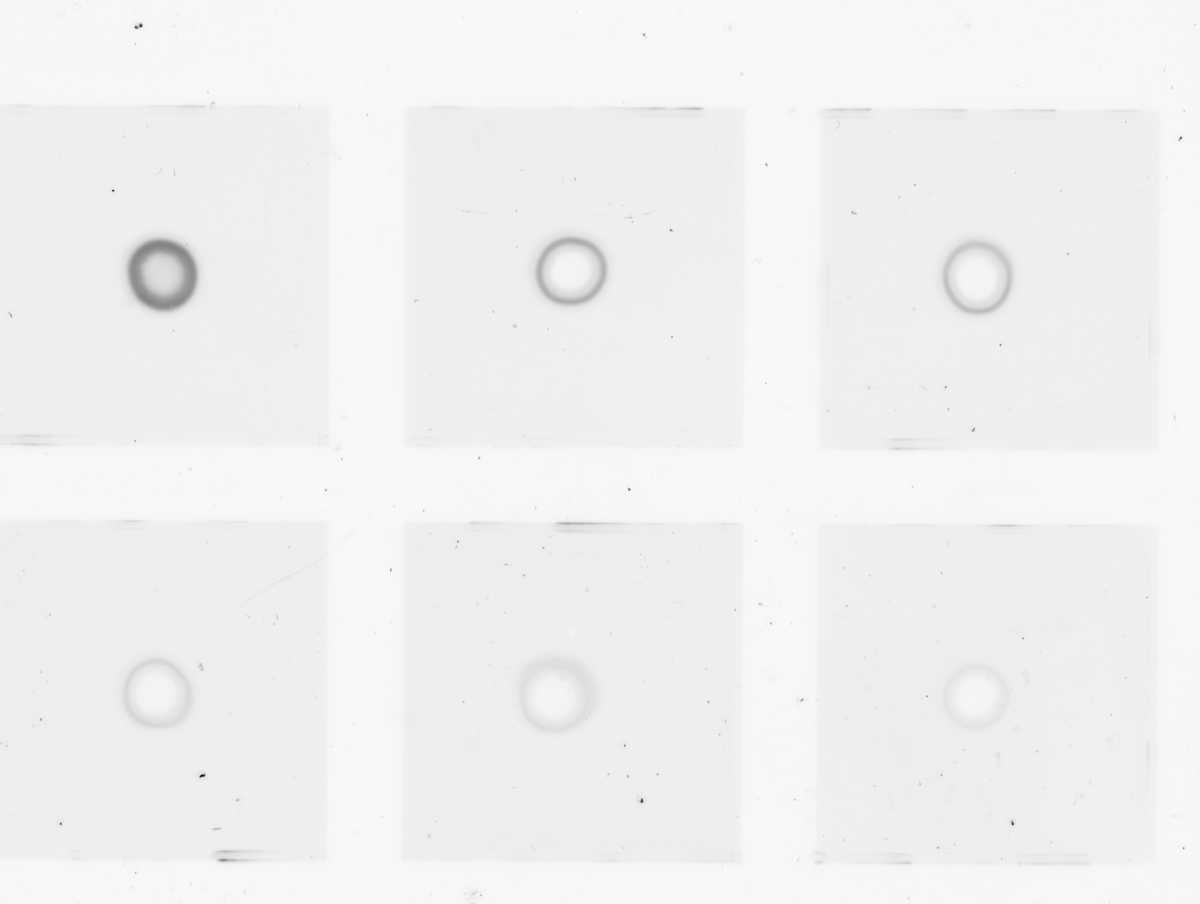

### Supplementary Video 2

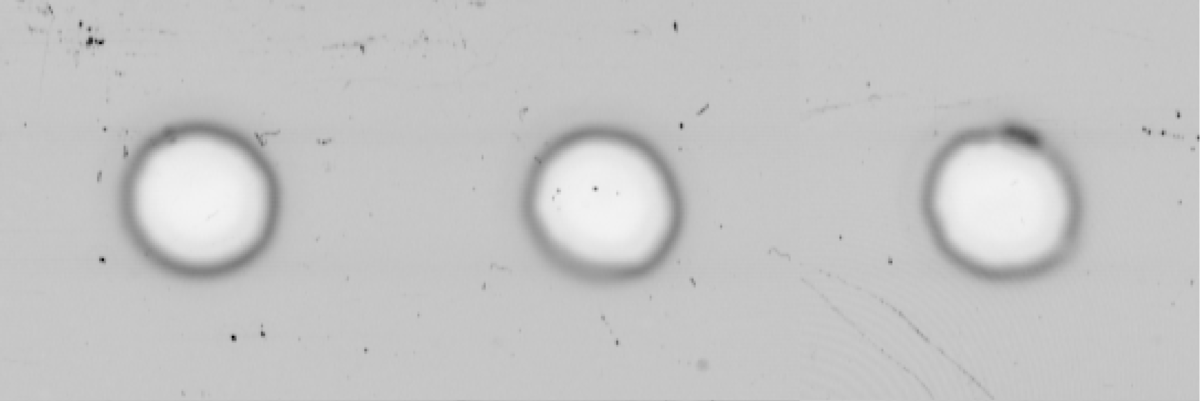

### Supplementary Video 3

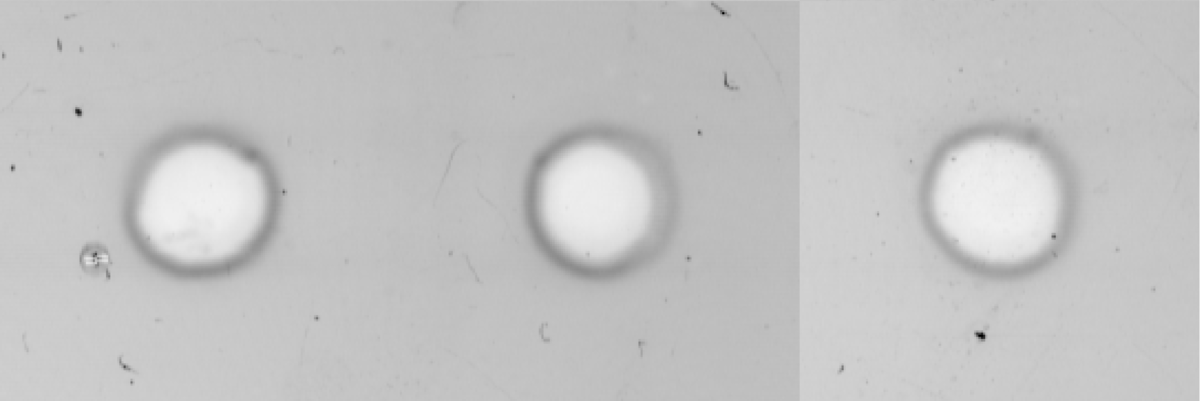

### Supplementary Video 4

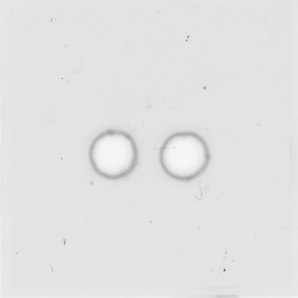

### Supplementary Video 5

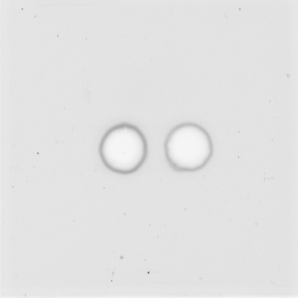

### Supplementary Video 6

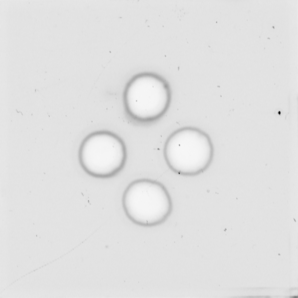

### Supplementary Video 7

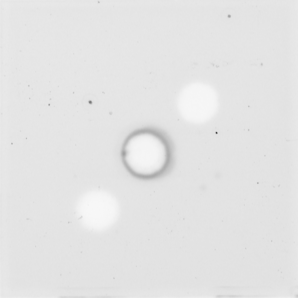

### Supplementary Video 8

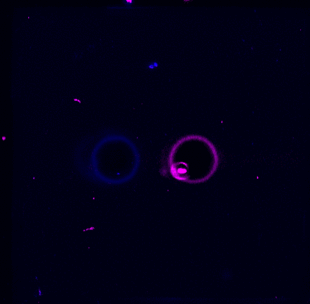

### Supplementary Video 9

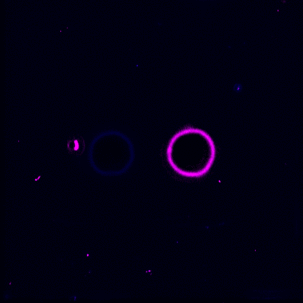
